# Histone divergence in *Trypanosoma brucei* results in unique alterations to nucleosome structure

**DOI:** 10.1101/2023.04.17.536592

**Authors:** Gauri Deák, Hannah Wapenaar, Gorka Sandoval, Ruofan Chen, Mark R. D. Taylor, Hayden Burdett, James A. Watson, Maarten W. Tuijtel, Shaun Webb, Marcus D. Wilson

## Abstract

Eukaryotes have a multitude of diverse mechanisms for organising and using their genomes, but the histones that make up chromatin are highly conserved. Unusually, histones from kinetoplastids are highly divergent. The structural and functional consequences of this variation are unknown. Here, we have biochemically and structurally characterised nucleosome core particles (NCPs) from the kinetoplastid parasite *Trypanosoma brucei*. A structure of the *T. brucei* NCP reveals that global histone architecture is conserved, but specific sequence alterations lead to distinct DNA and protein interaction interfaces. The *T. brucei* NCP is unstable and has weakened overall DNA binding. However, dramatic changes at the H2A-H2B interface introduce local reinforcement of DNA contacts. The *T. brucei* acidic patch has altered topology and is refractory to known binders, indicating that the nature of chromatin interactions in *T. brucei* may be unique. Overall, our results provide a detailed molecular basis for understanding evolutionary divergence in chromatin structure.

## Introduction

Nucleosomes are the basic unit of chromatin and control chromatin-associated processes in eukaryotic genomes. The nucleosome core particle (NCP) is composed of DNA wrapped around an octamer of four histone proteins (H2A, H2B, H3, and H4) and serves as a DNA compaction unit^1, 2^, inhibitor of transcription^3^, and dynamic molecular interaction platform^4^. In line with their role as architectural proteins, histone sequences are highly conserved, especially in well-studied eukaryotes^5, 6^. NCP structures from vertebrates^7, 8^, invertebrates^9^, and unicellular eukaryotes such as yeasts^10, 11^ have revealed remarkable conservation in global histone architecture and key sites of histone-DNA interactions. However, even small changes to histone primary sequences can have large structural and functional consequences on NCP structure^8, 12–14^.

Histone sequence variation and nucleosome structure in highly divergent eukaryotes are relatively understudied. The group Kinetoplastida is ranked amongst the most evolutionarily ancestral groups of parasitic protists and was estimated to have split from other eukaryotic lineages around 500 million years ago^15^. Kinetoplastida includes multiple pathogens, particularly those belonging to the *Trypanosoma* and *Leishmania* species. Of these, *Trypanosoma brucei* is a major clinical target, causing both human and animal trypanosomiasis^16–18^. In *T. brucei,* chromatin accessibility has a direct effect on antigenic variation, a key immune evasion mechanism contributing to its pathogenicity^19^. Further research on understanding how the trypanosome genome is organised and responds to stimuli could therefore have direct clinical and economic benefits.

Trypanosome chromatin has a number of unusual features. The *T. brucei* genome is organised into 11 large megabase chromosomes, a small, varying number of intermediate-sized chromosomes, and ∼100 minichromosomes^20^. Unlike in Metazoa, mitotic chromosome compaction levels in *T. brucei* are low (∼1.2-fold)^21^. Most genes are arranged in intron-less polycistronic transcription units^20^ and the chromatin context of transcription initiation and termination is in part defined by histones^22–24^. *T. brucei* histones have been found to be highly divergent^25^, and novel trypanosome-specific histone post-translational modifications^24, 26^ and chromatin interactors^27–30^ have been identified. However, the molecular details of how local nucleosome-level chromatin structure and molecular pathways in the nucleus intersect have been largely unexplored.

Here, we present the cryo-EM structure of the *T. brucei* NCP. The structure reveals altered histone-DNA contact sites, histone-histone interactions, and exposed histone surfaces compared to well-studied model eukaryotes. Globally, *T. brucei* NCPs are unstable and have reduced DNA binding. However, this instability is partly compensated by dramatic alterations in the electrostatic properties of *T. brucei* NCPs at the H2A-H2B binding interface. Furthermore, the surface topology, charge distribution, and binding properties of the *T. brucei* acidic patch are altered. Our phylogenetic analysis of kinetoplastid histones and their predicted structures reveals that these differences identified for *T. brucei* NCPs are likely conserved across the kinetoplastids. Overall, our study provides a molecular basis for understanding and further exploring how DNA compaction and chromatin interactions occur in *T. brucei*.

## Results

### Kinetoplastid histones are highly divergent

Previous reports have shown that a subset of trypanosomatid histones are highly divergent compared to model eukaryotes^25^. We performed an extended analysis of histone sequences from 22 distinct kinetoplastid genomes including multiple clades, such as those from the *Trypanosoma, Leishmania, Endotrypanum, Crithidia, Angomonas,* and *Perkinsela* species^31^. This analysis revealed a clear evolutionary divide between the kinetoplastids and a wide sample of eukaryotic taxa (Supplementary Fig. 1a). The divide was apparent across all histones, particularly H2A and H2B (Fig.1a). Within the kinetoplastids, our analysis pointed to multiple sub-groups of conserved sequences including segregation between the *Trypanosoma* and *Leishmania spp.* (Supplementary Fig. 1a). Aligned to *Homo sapiens* histones, the sequence identities of *T. brucei* histones was low, ranging from 40 to 60% (Fig.1a). Low conservation was apparent even when the predicted unstructured (and more commonly divergent) N- and C-terminal tails were excluded (Fig.1a). This prompted us to investigate whether histone sequence divergence in *T. brucei* leads to functional differences in nucleosome structure, assembly, and function.

### Reconstituting the *T. brucei* Nucleosome Core Particle (NCP)

To understand the basic unit of chromatin in *T. brucei,* we used an *in vitro* reconstitution approach to assemble recombinant *T. brucei* nucleosome core particles (NCPs). The four core histones were expressed and purified from *Escherichia coli* (Fig 1b & Supplementary Fig. 2a), refolded into octamers, and wrapped with the strong positioning Widom-601 DNA sequence using standard salt dialysis protocols^32–34^. The process of wrapping octamers with DNA required optimisation due to the presence of a soluble higher molecular weight species (Supplementary Fig. 2b-d, see Materials and Methods). We found that the thermal stability of *T. brucei* NCPs was significantly reduced compared to that of *H. sapiens* NCPs (Fig. 1c). Wrapping *T. brucei* histone octamers with 147 bp alpha-satellite DNA, another ‘strong’ positioning sequence^7^, also led to more unstable NCPs (Supplementary Fig. 2e).

**Figure 1:**
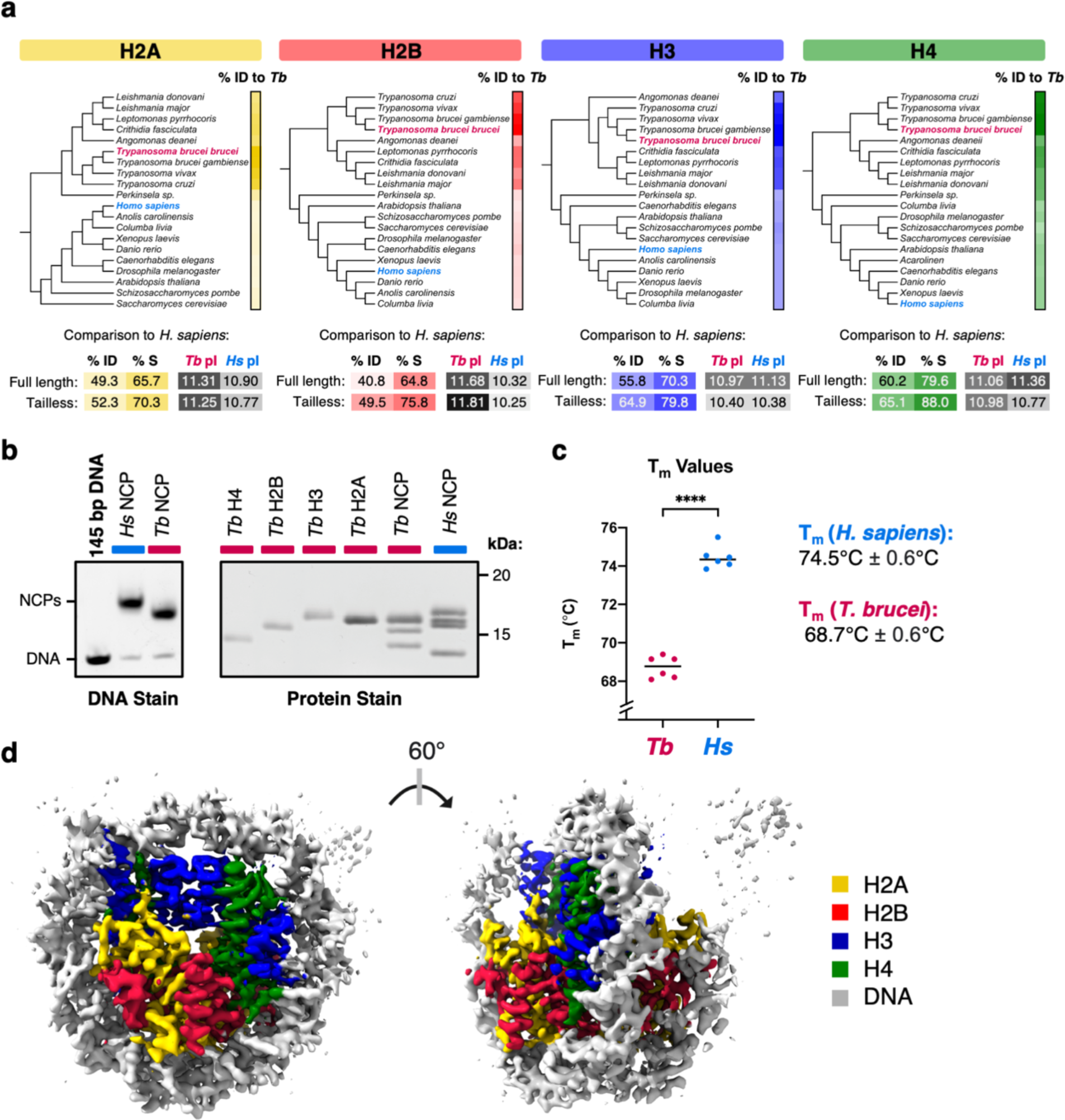
The *T. brucei* nucleosome core particle is evolutionarily divergent. **a**. Phylogenetic trees constructed from histone sequences of selected model organisms and kinetoplastid species. Scale bars on the right indicate pairwise percentage identity (%ID) to *T. brucei* colored from light to dark (range 40-100%). A comparison of *T. brucei* and *H. sapiens* histone sequences in terms of %ID, percentage similarity (%S), and predicted isoelectric points (pI) is shown below. **b.** Polyacrylamide gels of *in-vitro* reconstituted *T. brucei* and *H. sapiens* NCPs or component histones in their native (DNA stain) and denatured (protein stain + SDS) states. **c.** Mean melting temperatures (T_m_) of *T. brucei* and *H. sapiens* NCPs from three independent experiments (T_m_ values were calculated from the first derivative of melting curves shown in Fig. 3). **d**. 3.3 Å cryo-EM density map of the *T. brucei* NCP colored according to density attributed to histones and DNA.

On a coarse level, *H. sapiens* and *T. brucei* NCPs appeared structurally similar. By negative stain transmission electron microscopy, the *T. brucei* NCPs appeared as ∼10 nm disk shapes, reminiscent of NCPs from other species (Supplementary Fig. 3a). Furthermore, similar dimensions for *T. brucei* and *H. sapiens* NCPs were obtained by in solution small angle X-ray scattering (SAXS)(Supplementary Fig. 3b), suggesting that despite its differences in stability, at low resolutions, *T. brucei* nucleosome structure is maintained.

### The cryo-EM structure of *T. brucei* NCP reveals compressed histone architecture

In order to investigate the molecular details of divergence in *T. brucei* nucleosomes, we determined the structure of *T. brucei* NCP at a global resolution of 3.3 Å by single particle cryogenic electron microscopy (cryo-EM) (Fig. 1d, Supplementary Fig 3, Supplementary Movie 1). The low stability of the NCPs required us to perform mild chemical crosslinking prior to purification and sample preparation (Supp. Fig. 3c). The resolution was sufficient to allow us to build a model into the EM density for the core of the NCP (Fig.2a, Supplementary Fig. 4a, Table 1).

**Table 1:** Data collection parameters, processing information, and model validation metrics from cryo-EM structure determination of the T. brucei NCP

| Parameter | *T. brucei* NCP |
| --- | --- |
| **Data Collection** |  |
| Microscope | TFS Titan Krios |
| Detector | Gatan K3 |
| Acceleration voltage (kV) | 300 |
| Number of micrographs | 4913 |
| Frames per micrographs | 40 |
| Exposure time (s) | 7 |
| Dose per frame (e-/Å<sup>2</sup>) | 1.14 |
| Accumulated dose (e-/Å<sup>2</sup>) | 45.7 |
| Defocus range (- μm) | 1.5-3.2 |
| **Frames** |  |
| Alignment software | MotionCor2 |
| Frames used in final reconstruction | 2-40 |
| Dose weighting | Yes |
| **CTF** |  |
| Fitting software | CryoSPARC Patch |
| Correction | full |
| **Particles** |  |
| Picking software | CryoSPARC |
| Picked | 1,617,236 |
| Used in final reconstruction | 306,475 |
| **Alignment** |  |
| Alignment software | CryoSPARC |
| Initial reference map | Ab initio |
| low pass filter limit (Å) | 40 |
| number of iterations | 9 |
| local frame drift correction | no |
| **Reconstruction** |  |
| Reconstruction software | CryoSPARC |
| Box Size | 316 |
| Voxel size (Å) | 0.829 |
| Symmetry | C1 |
| Resolution limit (Å) | 1.658 |
| Resolution estimate (Å) | 3.28 |
| Masking | Yes |
| Sharpening (Å<sup>2</sup>) | -168 |
| EMDB ID | EMD-16777 |
| **Model Building** |  |
| Number of protein residues | 708 |
| Number of DNA residues | 254 |
| Bond length outliers | 0 out of 5593 |
| Bond angle outliers | 1 out of 7513 |
| Bonds (R.M.S.D) | 0.004 |
| Angles (R.M.S.D) | 0.571 |
| CaBLAM outliers (%) | 0.44 |
| Ramachandran favoured/allowed/outlier | 98.27/1.73/0.00 |
| Rotamer outliers (%) | 0.34% |
| Clash score | 8.77 |
| Model vs Data CC (mask) | 0.82 |
| EMringer | 1.34 |
| Fsc model vs map 0.5 | 0.261 (unmasked) |
| Molprobit score | 1.29 |
| PDB ID | 8COM |

The *T. brucei* NCP forms a characteristic coin-like shape, compacting and bending DNA around the core of the histone octamer (Fig. 1d). Although the histone secondary structure is largely preserved (Supplementary Fig. 4b), altered histone-DNA and histone-histone contacts lead to subtle changes in its overall architecture (Supplementary Fig. 5a). Compared to the *H. sapiens* NCP, the coin face of the *T. brucei* NCP exhibits compression along the horizontal axis (Fig. 2a; Supplementary Movie 2), with inwards shifts of histone-DNA interactions at super helical locations (SHL) 2 and 6 (Fig. 2b). We did not observe major uneven particle distribution in the cryo-EM map (Supplementary Fig. 3h & 3i), suggesting this shape was not due to directional anisotropy. The shift at SHL6 is coincident with a 2-residue insertion in Loop 2 of histone H2A. Although not conserved in sequence, this insertion is found across the *Trypanosoma spp.* (Supplementary Fig. 1b, Supplementary Movie 2). The longer length of this loop likely contributes to a ∼5 Å inward tilt of the ⍺2-helix of H2A (Fig. 2b). Higher flexibility of this region can also be observed from poorer local resolution (Supplementary Fig. 5b).

**Figure 2:**
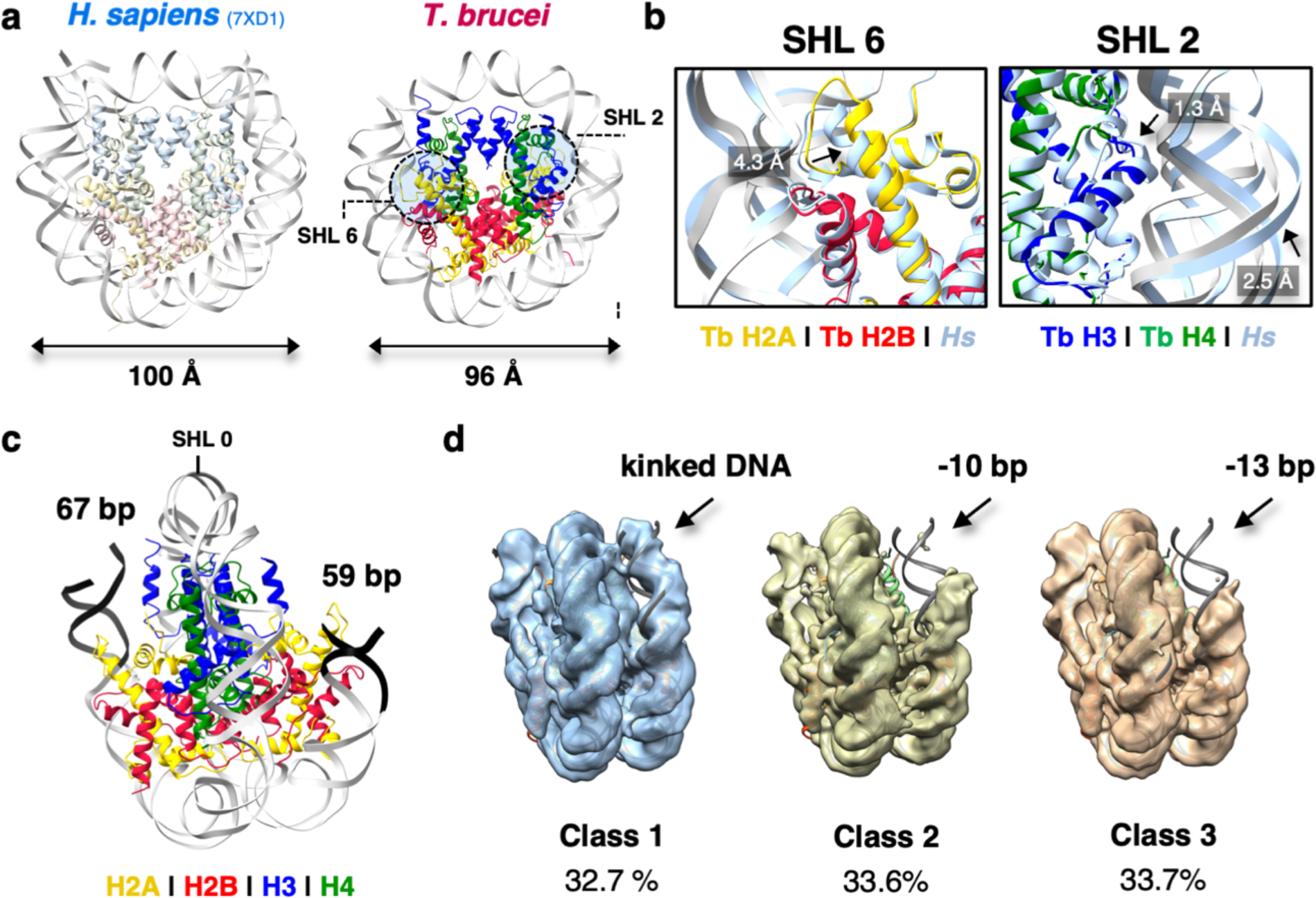
Histone morphology of the *T. brucei* NCP alters its interactions with DNA. **a.** Estimated diameters of *H. sapiens*^42^ and *T. brucei* NCP models from averaged measurements in angstrom (Å) between the phosphate atom of nucleotide -19 and nucleotide -59 in each DNA strand using ChimeraX^77^. **b.** Magnified view of the horizontal compression of the *T. brucei* NCP at SHL6 and SHL2. **c.** Poorer ordering and asymmetry of DNA ends (colored black) in the *T. brucei* NCP. **d.** Distinct 3D classes from cryo-EM processing showing different DNA conformations of the *T. brucei* NCP (Class 1 = 45,974 particles, Class 2 = 47,192 particles, and Class 3 = 47,429 particles).

At SHL2, we observed tighter packing of the ⍺1-L1 region of histone H3, termed the ‘H3 elbow’^4^. This likely occurs due to the loss of a bulky aromatic residue (*H. sapiens* (*‘Hs*’) Phe78 vs. *T. brucei* (*‘Tb*’) Gln75) that shifts the ⍺1-helix of H3 towards H4, increasing DNA bending in order to maintain arginine-phosphate interactions (Fig. 2b). This region is also more flexible (Supplementary Fig. 5b). Combined, these variations in histone architecture result in an oval-shaped NCP with altered DNA binding.

### The *T. brucei* NCP has more flexible entry/exit DNA

The ends of the wrapped DNA at the entry/exit site of the *T. brucei* NCP are poorly ordered in the cryo-EM density (Fig. 1c). We could reliably model only 126 bp of the 145 bp present, presuming that the missing ends of the DNA are flexibly tethered (Fig. 2c). Indeed, during image processing, distinct 3D classes of DNA-ordered states were obtained. Roughly one third of the data indicated a fully wrapped conformation, albeit with a bulged form of DNA, that was previously observed in *Xenopus laevis* NCPs as a precursor to unwrapping^35^.The remaining two thirds lacked density around the DNA entry/exit sites (Fig. 2d), indicating flexibility of DNA ends.

The pseudo-symmetry in the *T. brucei* NCP is broken, with one end of the DNA being considerably more disordered. This is likely due to the asymmetry of the Widom 601 sequence^36, 37^ and is consistent with partial asymmetric unwrapping observed previously^35, 37–40^. The inwards compression of DNA observed at SHL2 and 6 and the more oval shape of the *T. brucei* NCP (Fig. 2b) may contribute to this splaying of DNA ends. Despite DNA entry/exit site flexibility, we note that the histone-DNA register at the core of the histone octamer is maintained overall, both from comparison to other structures and a hydroxyl radical footprinting assay (Supplementary Fig. 5c).

Unsurprisingly, the density for *T. brucei* histone tails was low, arising from a high degree of disorder^41^. However, compared to *H. sapiens* and *X. laevis* EM density maps at similar resolutions^35, 42^, there is poorer ordering for the C-terminal tail of H2A and the N-terminal tail of H3, which engage the final ∼13bp of straight entry/exit DNA on the NCP^43^. *T. brucei* has a number of amino acid substitutions in histones H2A and H3 in these terminal regions (Supplementary Fig. 4b). Notably, *Hs* Arg53 is altered to *Tb* Gln50 in the ⍺N helix of histone H3 (Supplementary Fig. 5d-e). Substitutions at this position were previously shown to be critical for destabilization of entry/exit DNA binding in nucleosomes containing human H3 variants^14, 44^. DNA end accessibility was also observed in *T. brucei* NCPs using an exonuclease assay (Supplementary Fig. 5f). Overall, the oval morphology and poor ordering of H3/H2A tails likely leads to flexibility of entry/exit DNA in the *T. brucei* NCP.

### Alterations in histone-histone interfaces in *T. brucei* NCP lead to instability

Despite the overall conservation of the fold of the *T. brucei* histone octamer, notable changes are present at histone-histone interfaces in the *T. brucei* NCP. Within the H3-H4 tetramer, amino acid substitutions affecting steric packing between H3-H3 and the H3-H4 interface likely alter the stability of the *T. brucei* tetramer and octamer. Electrostatic interactions and the hydrophobic core of the H3-H3 four-helix bundle are disrupted (*Hs* H3-Ala111, *Hs* H3-Ala114, and H3-Leu109 to *Tb* H3-Cys108, *Tb* H3-Ser111, and *Tb* H3-Arg106, respectively) (Fig. 3a). Single substitutions in this region have previously been shown to destabilize NCPs^12, 45^. At the H3-H4 interface, altered hydrophobic interactions arise due to the substitution of *Hs* H3-Phe104 to *Tb* H3-Leu101 (Supplementary Fig. 6g). Interestingly, substitutions at this position were also reported in unstable nucleosomes^13, 46^. Multiple sites that are critical in imparting stability to the H3-H4 tetramer are therefore altered in *T. brucei*.

**Figure 3:**
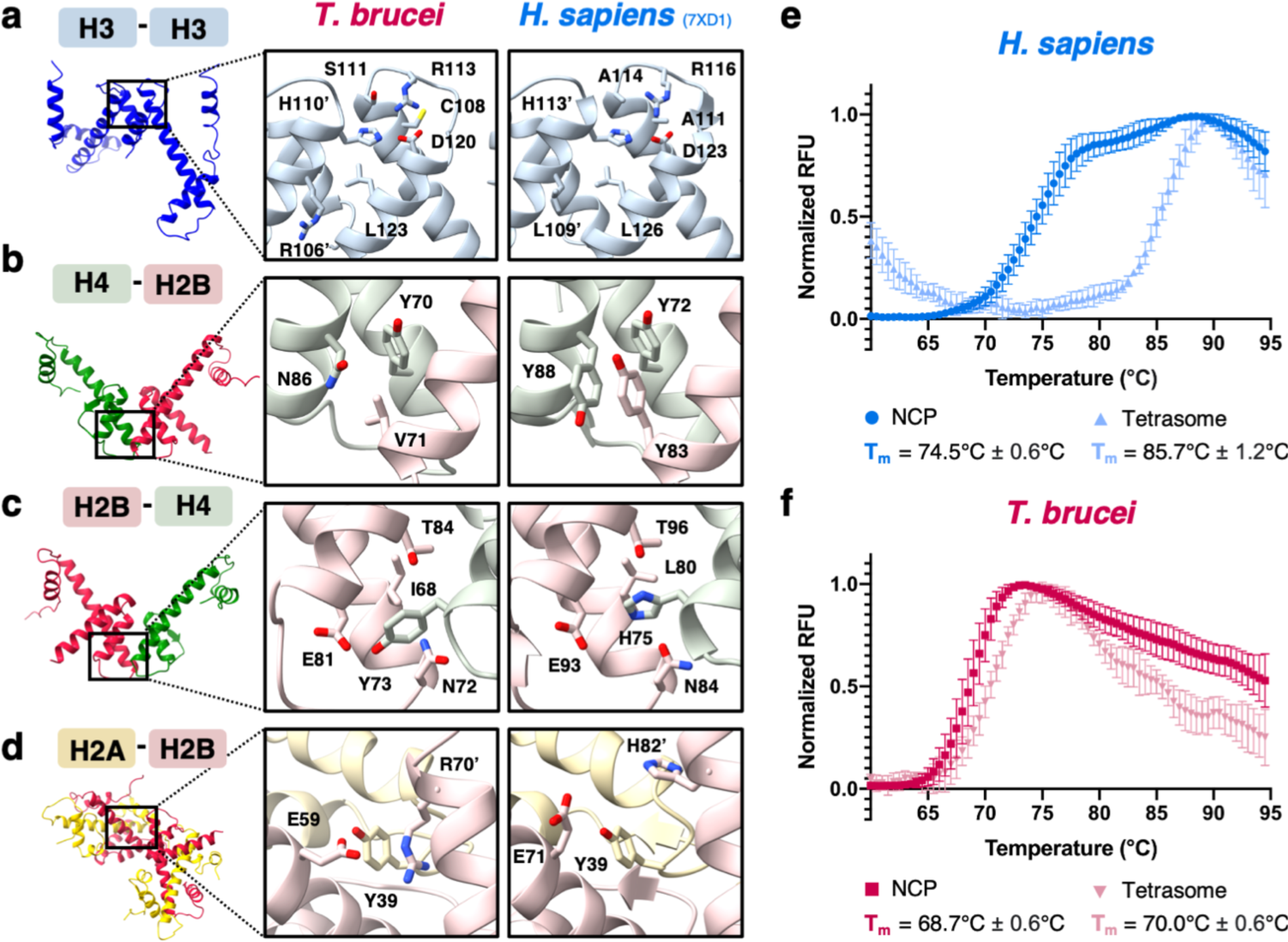
The *T. brucei* NCP is less stable and has a monophasic disassembly pathway. **a-d.** Magnified views comparing key amino acid interactions at histone-histone interfaces in the *T. brucei* and *H. sapiens* NCPs^42^. Apostrophes indicate the second copy of a histone if two histones of the same type are present. **e.** Biphasic thermal denaturation curves of *H. sapiens* NCPs (darker blue) and monophasic denaturation curves of *H. sapiens* H3-H4 tetrasomes (lighter blue) *(n* = 6). **f.** Monophasic thermal denaturation curves of *T. brucei* NCPs (darker pink) and *T. brucei* H3-H4 tetrasomes (lighter pink) *(n* = 6). Data points in **e.** and **f.** are normalized average values ± SD.

Intriguingly, the H3-Cys110 pair that has been implicated in redox sensing^47^ is not structurally conserved. Instead, it is replaced by two cysteine pairs that are spatially separated (*Tb* H3-Cys108, *Tb* H3-Cys126).

At the dimer-tetramer interface, the *T. brucei* H2B-H4 four-helix bundle exhibits a loss of aromatic and hydrogen bonding interactions. A π-stacking network mediated by *Hs* H2B-Tyr83, H4-Tyr72, and H4-Tyr88 is absent due to the loss of two of the three tyrosines in *T. brucei* (Fig. 3b). A similar loss has been documented in the unstable nucleosome-like structures formed by Melbournevirus histone doublets^48^. Additionally, a multivalent hydrogen bonding network centred on H2B-His75 in *H. sapiens* NCPs is disrupted through substitution to H2B Tyr-73 in *T. brucei* (Fig. 3c). Conversely, a reinforcement of binding likely occurs at the H2A-H2B dimer-dimer interface, where *Tb* H2A-Tyr39 and *Tb* H2B’-Arg70 likely form a cation-π interaction that is lacking in *H. sapiens* (Fig. 3d).

These observations agree well with our attempts to reconstitute hybrid *T. brucei/ H. sapiens* histone octamers. Octamers with *Hs* H3 + *Tb* H2A, H2B, and H4 could assemble due to the compatibility of *Hs* H3-*Hs* H3 and minor changes at the H3-H4 interface described (Supplementary Fig. 2d; Supplementary Fig. 6f). NCPs reconstituted with this octamer had slightly higher thermal stability (Supplementary Fig. 6h), indicating that the *Tb* H3-H3 interface has lower cohesion. However, more extensively altered histone-histone interfaces prevented the assembly of hybrid octamers containing *Hs* H2A + *Tb* H2B, H3, and H4 (Supplementary Fig. 6d) or *Hs* H2A-H2B + *Tb* H3-H4 (Supplementary Fig. 6e). Although heterodimerization of *Hs* H2A and *Tb* H2B itself was possible (Supplementary Fig. 6d), the formation of a stable H2A-H2B dimer-dimer interface in the context of an assembled octamer was prevented.

The altered protein-protein interactions in *T. brucei* NCPs drive not only lower thermal stability but also result in an altered NCP disassembly pathway (Fig. 3e-f). As previously reported^49, 50^, *H. sapiens* NCPs exhibit biphasic disassembly, whereby the first step involves H2A-H2B disengagement from the H3-H4 tetramer followed by breakdown of the tetramer itself (Fig 3e). Unusually, *T. brucei* NCP disassembly is a single event (Fig.3f). The major drivers are probably the instability of the dimer-tetramer interface and the instability of the tetramer itself, causing both NCP disassembly at a lower temperature and the disassembly of reconstituted *T. brucei* H3-H4 tetrasomes at a similar temperature to *Tb* NCPs (Fig. 3e). The majority of the amino acid substitutions outlined above are conserved in the *Trypanosoma spp.* and would be expected to produce comparable effects in other *Trypanosoma* NCPs (Supplementary Fig. 1b).

### The *T. brucei* NCP has reduced overall protein-DNA interactions

Alterations in histone-DNA contacts also contribute to the instability of *T. brucei* NCPs. Similarly to *H. sapiens* NCPs, the structure of the histone folds maintains the global alignment of helix dipoles to the DNA phosphate backbone. The total number of histone-DNA hydrogen bonding interactions in the two NCPs is near-equivalent (Fig. 4a). However, the distribution of these bonds is altered across the four histone types (Fig. 4a) and changes in electrostatic interactions at specific DNA contact points lead to an overall reduction of DNA binding in the *T. brucei* NCP.

**Figure 4:**
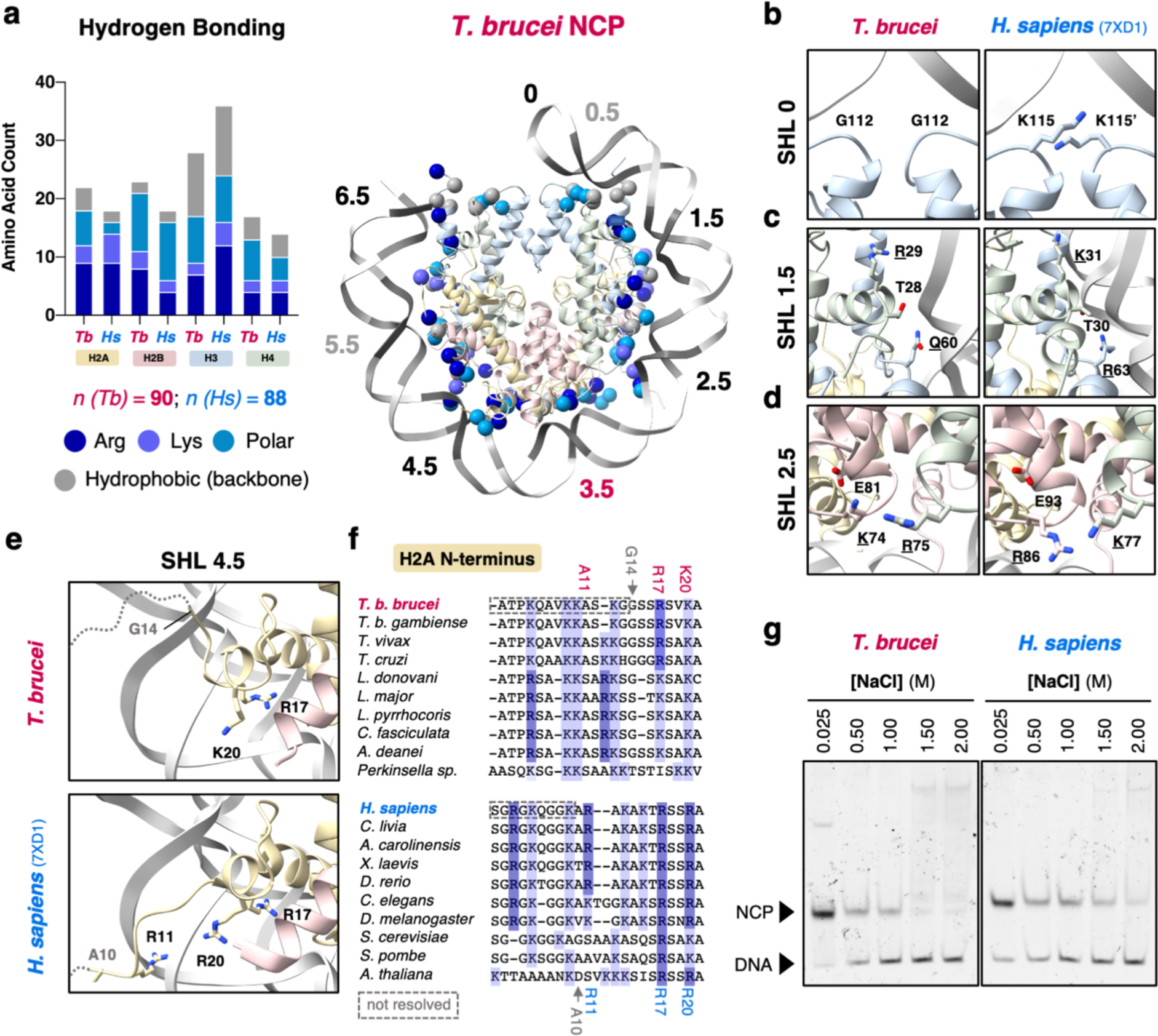
Protein-DNA contacts in the *T. brucei* NCP have a different distribution and result in weaker DNA binding. **a.** Frequencies of different amino acids forming hydrogen bonds with DNA in the *T. brucei* and *H. sapiens* (PDB: 7XD1)^42^ NCPs predicted by ePISA^78^ are shown on the left. The total number of interacting residues for each NCP (n) is stated below. On the right, the DNA contacting residues predicted for *T. brucei* are mapped onto the model of the NCP as spheres. SHL annotations are colored according to reduced (black), increased (pink), or similar (grey) interactions with the DNA phosphate backbone. **b.-e.** Reduced protein-DNA interactions at SHL0, 1.5, 2.5, and 4.5 in the *T. brucei* NCP compared to the *H. sapiens* NCP. **f.** Multiple sequence alignment of the N-terminal tail of histone H2A at SHL 4.5, with DNA-contacting residues from part **e.** highlighted. **g.** Native polyacrylamide gels stained for DNA showing DNA unwrapping in *T. brucei* and *H. sapiens* NCPs after incubation at different NaCl concentrations for 1h on ice.

At SHL 0 (the dyad), two lysines from histone H3 (*Hs* Lys115) are substituted for glycine (*Tb* Gly112) (Fig. 4b). At SHL1.5, a critical arginine in histone H3 (*Hs* Arg63)^7^ is substituted for *Tb* Gln60 (Fig. 4c). Interestingly, SHL2.5 features a case of histone co-evolution, where histones H2B and H4 in *T. brucei* have a spatially coincident arginine-to-lysine and lysine-to-arginine substitutions respectively (Fig. 4d). This prevents steric clash compared to a single substitution and allows for extended interactions with *Tb* H2B-Glu81. However, the new interaction occurs at the expense of the proximity of *Tb* H2B-Lys74 to DNA (Fig. 4d).

The N-terminal tail of H2A, which typically straddles the minor groove at SHL4.5^30, 39, 47, 48^, is poorly ordered in *T. brucei* (Fig. 4e) and a phosphate-interacting residue, *Hs* Arg11, is replaced by *Tb* Ala11 (Fig. 4e-f). The C-terminal helix of histone H2B is displaced and is shorter in length by two residues. The lysine that normally anchors the helix to DNA is therefore lacking (Supplementary Fig. 7a). These changes in H2A and H2B are conserved across the kinetoplastid species (Fig. 4f, Supplementary Fig. 7b). Cumulatively, the loss of DNA contacts described above suggest weaker DNA binding by the *T. brucei* NCP and this is supported by both lower thermal stability (Fig. 1c) and lower resistance to increasing salt conditions compared to the *H. sapiens* NCP (Fig. 4g).

### A concentrated cluster of positive charge drives DNA binding by H2A-H2B at SHL3.5

While overall DNA-protein contacts are reduced in the *T. brucei* NCP (Fig. 4, Supplementary Fig. 5d), a unique cluster of positively charged residues reinforces DNA binding at a discreet and unusual position on the NCP: the dimer-dimer interface of histones H2A and H2B at SHL3.5. Compared to *H. sapiens, T. brucei* H2A and H2B have higher calculated net positive charge and this holds true even when the disordered histone tails are discounted (Fig. 1a). The structure shows higher electrostatic surface potential at SHL3.5 (Fig. 5a) and the interface features net gain of positively-charged side chains from two charge swapping events (*Hs* H2A-Glu41 to *Tb* H2A-Arg41, *Hs* H2B-Glu35 to *Tb* H2B-Arg23), two additional arginine residues (*Tb-*H2B Arg70, *Tb-*H2B Arg75), and a lysine-to-arginine substitution (*Hs* H2A-Lys36 to *Tb*-H2A-Arg36; Fig. 5b;). The dense cluster of positive charge dramatically alters the overall predicted dipole moment of *T. brucei* and *H. sapiens* histone octamers, where the direction of the dipole is oriented opposite to the dyad at SHL 3.5 (Fig. 5c). Interestingly, this may be a kinetoplastid-specific adaptation based on conservation at the sequence level (Supplementary Fig. 8a). Furthermore, this interface is similarly charged in models of histone octamers generated using AlphaFold2^51^ from five representative kinetoplastid species, but not other NCP structures (Supplementary Fig. 8b).

**Figure 5:**
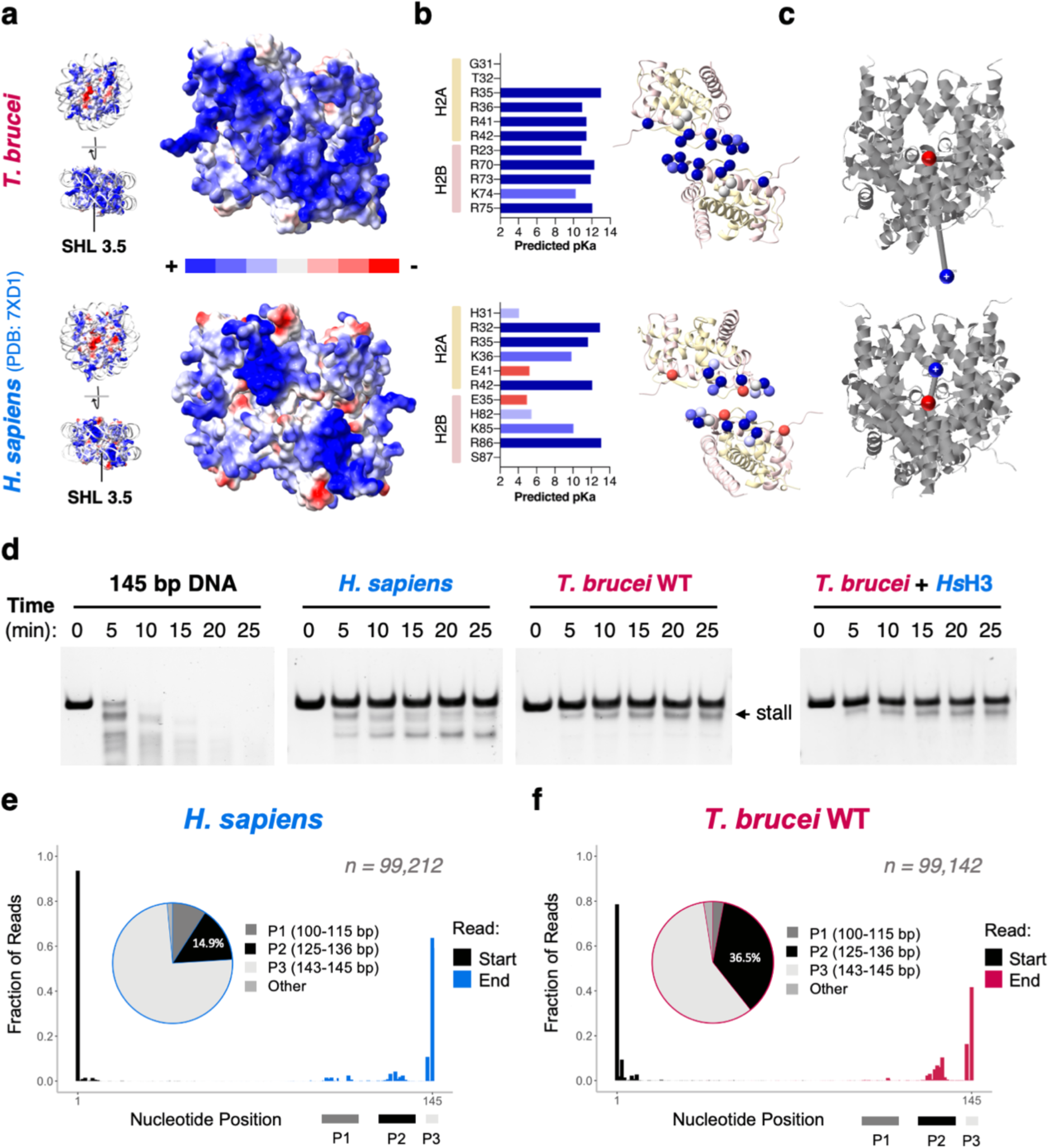
A unique mode of DNA binding at SHL 3.5 in the *T. brucei* NCP. **a.** Surface electrostatics showing that *T. brucei* histone octamers have a higher density of positive charge at SHL 3.5 compared to *H. sapiens*^42^. **b.** On the left, predicted pKa values^79^ of residues contributing to the electrostatic potential at SHL 3.5 in *T. brucei* and *H. sapiens* NCPs. On the right, the same residues shown as spheres mapped onto the structures of the *T. brucei* and *H. sapiens* H2A-H2B dimers. **c.** Graphical representation of the predicted overall molecular dipole moments^80^ of *T. brucei* (∼1093 Debye) and *H. sapiens* (∼294 Debye) histone octamers (positive end = blue, negative end = red). **d.** Limited micrococcal nuclease digestion of DNA alone, *H. sapiens* NCPs, *T. brucei* NCPs (‘WT’), and chimeric *T. brucei* NCPs with *H. sapiens* histone H3 (‘*T. brucei* + *Hs*H3’). **e.** Start and end positions of sequenced *H. sapiens* MNase digestion products from **d.** with fractions of reads present in peaks P1, P2, and P3 displayed in a pie chart; **f.** The same analysis as in **e.** repeated for *T. brucei* MNase digestion products.

The observation of locally reinforced DNA interactions correlates well with micrococcal nuclease (MNase) digestion assays. Although the rate of MNase digestion of *T. brucei* NCPs was comparable to *H. sapiens* NCPs (Supplementary Fig. 8c), we observed a characteristic digestion band for *T.* brucei NCPs not observed in *H. sapiens* NCPs (Fig. 5d). We hypothesise this corresponds to the MNase pausing at a stall point, prior to recovering and continuing to digest the DNA. The same experiment repeated with chimeric NCPs with *H. sapiens* H3 and *T. brucei* H2A, H2B, and H4 (‘*T. brucei* + *Hs*H3’), did not change this pattern, suggesting the role of *T. brucei* H2A-H2B dimers in the stall (Fig. 5d). To approximate DNA length at the ‘stall point’, we prepared a custom DNA ladder, which showed that the stall point is roughly 10-14 bp shorter than full length DNA in *T. brucei* NCPs (Supplementary Fig. 8d).

We further mapped the stall point by sequencing both *T. brucei* and *H. sapiens* MNase reaction products (Fig. 5e-f). As expected, the digestion occurs preferentially from the more accessible flexible DNA end (Fig. 2c). The stall point corresponds to ∼12 bp digestion in *T. brucei* (highest data point in Peak 2, Fig. 5f) adjacent to where H2A-H2B first begin to contact DNA (Supplementary Fig. 8e). In contrast to *H. sapiens*, further digestion products can barely be detected (Peak 1 in Fig. 5e-f; Supplementary Fig. 8e) . This is in good agreement with the presence of the highly basic region in the *T.brucei* NCP at SHL 3.5 and may serve as a unique kinetoplastid mechanism to anchor DNA firmly to the base of the nucleosome despite unstable DNA binding at the dyad and at entry/exit sites.

### Altered octamer topology in the *T. brucei* NCP leads to an atypical acidic patch

The acidic patch is a negatively charged region formed between histones H2A and H2B that serves as an interaction platform for many chromatin-associated proteins^4^. It has also been implicated in the formation of higher order chromatin structure^7, 52^. Remarkably, given the high sequence divergence, the negative charge of the canonical acidic patch residues in the *T. brucei* are conserved (Fig. 6a). However, the coulombic surface potential of the *T. brucei* acidic patch exhibits stark differences to *H. sapiens* (Fig. 6b). The overall shape of the *T. brucei* acidic patch appears narrower and the estimated surface area of the patch is smaller (4498 Å^2^ vs. 4702 Å^2^; Fig. 6b). This is likely driven by the two-residue insertion in *Tb* H2A Loop 2 that allows an inwards shift of the ⍺2 helix of H2A at SHL6 (Fig. 2b).

**Figure 6:**
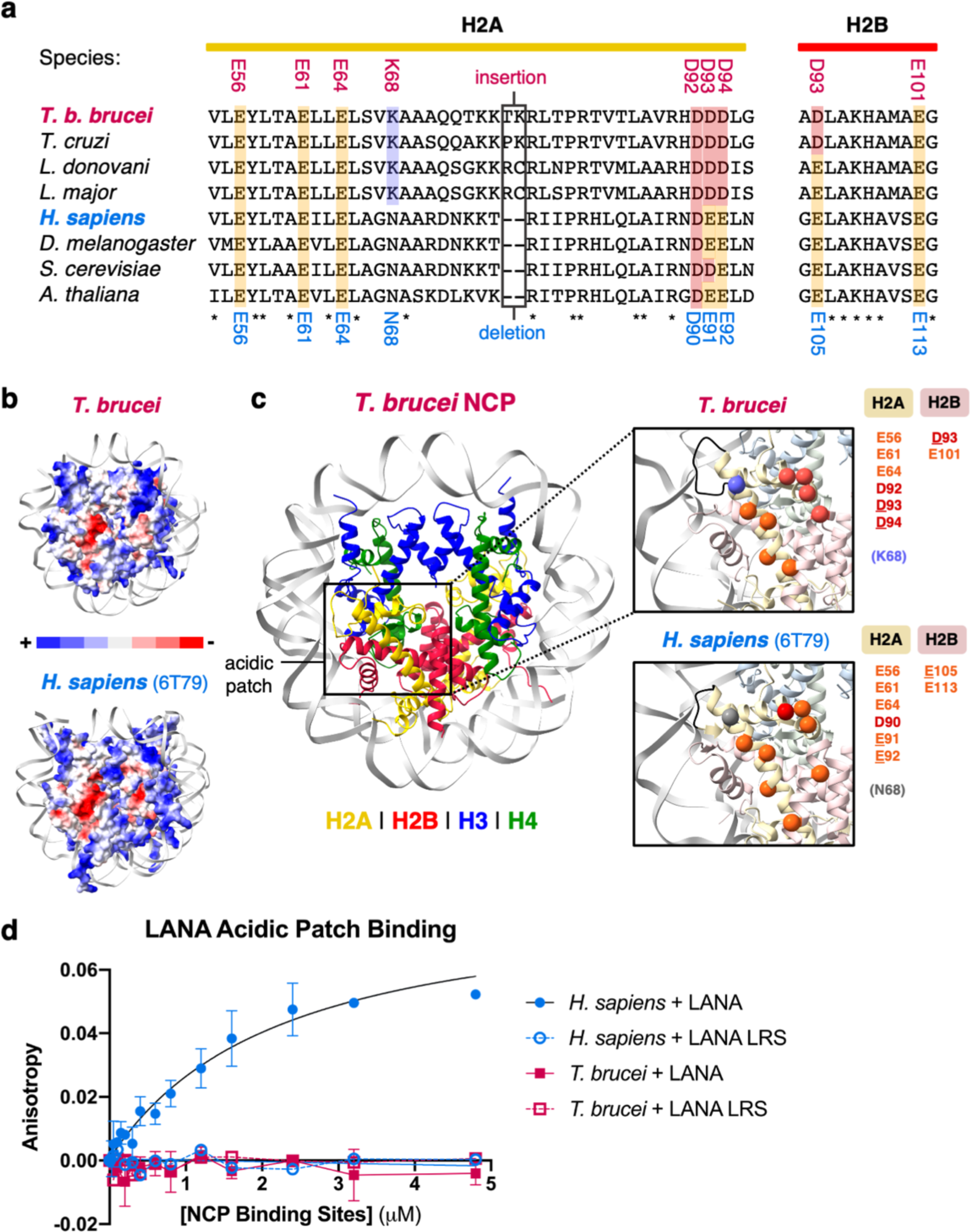
The *T. brucei* acidic patch is highly atypical and refractory to well-characterized binders. **a.** Multiple sequence alignments of acidic patch regions in H2A and H2B, where residues are highlighted and coloured Glu = orange, Asp = red, Lys = purple, Asn = grey. and the residue numbers for *T. brucei* (above) and *H. sapiens* (below) are indicated. **b.** The acidic patch region differs when visualizing the electrostatic surface representation of the *T. brucei* (top) and *H. sapiens* (PDB: 6T79^81^, bottom) NCPs. **c.** The *T. brucei* NCP with a magnified view of the acidic patch and compared to the *H. sapiens* acidic patch below. Acidic patch residues are shown as spheres and coloured using same scheme as **a.** the 2-residue insertion in Loop 2 of *T. brucei* H2A is indicated in black. **d.** Fluorescence polarization assay showing binding of a FITC-tagged LANA peptide and a non-binding mutated LANA peptide (L8A R9A S10A, ‘LANA LRS’) to *H. sapiens* and *T. brucei* NCPs (K_D_ for *H. sapiens* NCPs = ∼2.38 μM, others ND assuming two independent binding sites per NCP).

A number of other substitutions are present in this region, particularly in histone H2A (Fig. 6a). While three of the canonical residues have undergone mild glutamate-to-aspartate substitutions (*Tb* H2A-Asp93, *Tb* H2A-Asp94, and *Tb* H2B-Asp93), a lysine substitution (*Tb* H2A-Lys68 from *Hs* H2A-Asn68) is in close proximity to the patch (Fig. 6c). This substitution is conserved across the kinetoplastids (Supplementary Fig. 9a) Interestingly, mutation of *Hs* H2A-Asn68 was recently shown to reduce binding to multiple chromatin-associated proteins in a proteome-wide screen.^53^ Both the surface charge and shape alterations are also conserved in our predicted models of five other kinetoplastid histone octamers (Supplementary Fig. 9b).

These differences in the *T. brucei* acidic patch likely have direct functional consequences for chromatin reading. A prototypical acidic patch binder, the Kaposi’s Sarcoma Herpesvirus Latency Associated Nuclear Antigen (LANA) peptide^54^, binds well to the *H. sapiens* NCP but showed no detectable interaction to the *T. brucei* NCP in a fluorescence polarization assay (Fig. 6d). Similar results were obtained for an acidic patch binder with an altered binding mode (Supplementary Fig. 9c), the Prototype Foamy Virus GAG peptide (PFV-GAG)^55^, using both fluorescence polarization (Supplementary Fig. 9d) and electrophoretic mobility shift assays (Supplementary Fig. 9e). Discrete mutations to individual components of the *T. brucei* acidic patch were not sufficient to rescue the binding of PFV-GAG (Supplementary Fig. 9a and 9e). These results reveal extensive divergence of this common protein interacting hub and indicate that the kinetoplastids likely have altered chromatin-protein and higher order chromatin interactions.

## Discussion

In this study, we present the cryo-EM structure and biochemical characterisation of the nucleosome core particle from the parasitic protist *T. brucei*. Despite extensive divergence of *T. brucei* histone sequences (Fig. 1), the structure exhibits remarkable conservation of overall histone fold architecture. This is consistent with structures from other divergent organisms such as the parasite *Giardia lamblia*^56^, the archaeon *Methanothermus fervidus*^57^, or the *Marseilleviridae*^48, 58^ giant viruses. However, subtle differences in histone packing and specific histone amino acid substitutions give rise to properties unique to the *T. brucei* NCP. The *T. brucei* NCP has a compressed shape (Fig. 2, Supplementary Movie 2) and four out of seven DNA contact points at half-integral SHLs have lost critical histone-DNA interactions, leading to flexible DNA ends and low stability *in vitro* (Fig. 2-4). A striking compensation mechanism occurs via H2A-H2B dimers at SHL3.5, whereby DNA binding is increased by a concentrated cluster of basic residues (Fig. 5). Furthermore, kinetoplastid-specific alterations lead to an altered topology and charge of common protein interaction interface, the acidic patch (Fig. 6). By comparing the molecular differences in nucleosomes between trypanosomes and conventional eukaryotic systems, we can hope to understand both the evolutionary constraints and diversity that directs DNA-associated mechanisms.

Early studies of trypanosome chromatin showed that *T. brucei* chromatin is less stable^59^ and has reduced higher order compaction than is observed in model eukaryotes^21, 60, 61^. Our results indicate that chromatin instability in *T. brucei* is inherent at the mononucleosome level via weakened histone-histone interfaces, histone-DNA contacts, and flexible entry/exit DNA (Fig. 2-4). Chromatin arrays constructed from other nucleosomes with DNA end flexibility^46, 56, 62^ were also found to favour more open chromatin conformations, particularly due to alterations in inter-nucleosomal DNA path^62^. Future experiments on chromatin arrays would help explore if entry/exit DNA flexibility described here could explain the more open chromatin in *T. brucei*.

However, despite overall lower DNA binding, histones H2A and H2B in *T. brucei* were previously shown to be bound to DNA more persistently than H3 and H4^63^. This is consistent with our finding that H2A-H2B reinforce DNA binding at SHL3.5 via a large positive dipole (Fig. 5). The significance of tight DNA binding by H2A-H2B requires further investigation but we speculate that it may affect a variety of chromatin-based processes. For example, RNA polymerase II transcription is known to be stalled at defined points corresponding to DNA-histone contacts while transcribing a nucleosome template^64–67^. Transcription in *T. brucei* is expected to be highly atypical^68^ and future studies could reveal whether the altered DNA binding properties of their nucleosomes affect this and other processes.

Recent work has shown the complexity of histone post-translational marks in *T. brucei*^24, 26, 27^ and *T. cruzi*^69, 70^. Although most of the N- and C-terminal histone tails are not resolved in our structure, we could place 47 reported histone marks onto our structure^24, 26^ (Supplementary Fig. 10a). Intriguingly, the kinetoplastid-specific H2A-Lys68, which alters the local environment of the acidic patch (Fig. 6b), has been reported to be trimethylated (Supplementary Fig. 10a). Conceivably, this and other modifications may act as switches to alter chromatin binding properties. We also note a cluster of phosphorylation/acetylation sites at the N-terminal end of H2A (Supplementary Fig. 10a) that could serve to further destabilise DNA binding at SHL 4.5 (Fig. 4e). Beyond histone post translational modifications, histone variants in *T. brucei* play a key role in modulating transcription^22–24^ and controlling the parasite’s variant surface glycoprotein immune evasion system^19^. Further work comparing the canonical structure of *T. brucei* nucleosomes to variants would be of great interest to help explain the biology behind antigenic variation and transcription initiation/termination.

Of the four common interaction interfaces commonly co-opted by chromatin-binding proteins^4^, all display both sequence and structural variation in *T. brucei*. For example, the acidic patch in *T. brucei* has altered topology and is refractory to known binders (Fig. 6). Interestingly, the narrow topology of the patch accompanied by an insertion in H2A in *T. brucei* is inverse to the widening of the patch with an insertion in H2B in the nucleosome structure of the parasite *G. lamblia*^56^ (Supplementary Fig. 9a). The divergence of this interface suggests that trypanosome chromatin interactions may also have divergent properties. Since the acidic patch has been identified as a common interaction site in other species^4^, we expect that other identified factors in *T. brucei* involved in regulating histone post-translational modifications^27^, chromatin remodelling^28, 29^, antigenic variation^30^, or yet unidentified pathways may use this mechanism.

However, a mechanistic understanding for chromatin interactions in *T. brucei* is largely missing and our search for acidic-patch binders *in T. brucei* was challenging due to low conservation or poor annotation of potential homologs. Despite this, promising candidates could include *T. brucei* Dot1A and Dot1B, homologs of the Dot1 histone methyltransferase in higher eukaryotes^71–73^. The catalytic fold of *T. brucei* Dot1A/B seems to be conserved^34^ when modelled using AlphaFold2^51^ (Supplementary Fig. 10b), but the loop that engages the acidic patch in human Dot1L^73^ (Supplementary Fig. 10c) differs in both sequence and predicted structure in *T. brucei* (Supplementary Fig. 10d &e). This suggests that the local binding interactions to chromatin by *T. brucei* Dot1A/B may differ. It will be a fascinating avenue of study to probe the effects of divergence on chromatin recognition in kinetoplastid parasites.

Remarkably, our extensive phylogenetic analysis and modelling revealed that a majority of our findings are conserved within the kinetoplastids, including known pathogens (Supplementary Figure 1a, 8a-b, 9a-b). For example, conservation of the altered acidic patch includes clinically relevant targets from the *Trypanosoma* and *Leishmania* species. (Supplementary Fig. 9a-b). This opens a possible therapeutic avenue for targeting the acidic patch^74, 75^ allowing specific targeting of kinetoplastids over their human or animal hosts. A global chromatin disruption mechanism would have high utility for combating diseases such as animal trypanosomiasis where the genetic diversity of *Trypanosoma* species has hindered drug development^17^ and drug resistance is a current challenge^16, 76^.

## Data Availability

The cryo-EM density map and associated meta data for the *T. brucei* NCP have been deposited at the Electron Microscopy Data Bank under accession number EMD-16777. Raw micrographs have been uploaded to EMPIAR-XXX. The atomic coordinates of the *T. brucei* NCP have been deposited in the Protein Data Bank under accession number 8COM. The MNase sequencing results have been deposited to the Gene Expression Omnibus NCBI database under accession number GSE226029.

## Acknowledgements

We thank Robin Allshire, David Horn, Christopher Wood, Elizabeth Blackburn, and members of the Wilson lab for helpful discussions. We thank Kurt Davey for gift of the alpha satellite repeat plasmid and Christian Janzen for gift of the *T. brucei* histone plasmids. Human histones were supplied from Addgene from the Landry lab. We thank Logan Mackay in SIRCAMS, School of Chemistry, University of Edinburgh for mass spec analysis. We also thank the Technical Services Sequencing Facility at The Institute of Genetics and Cancer, University of Edinburgh for processing hydroxyl radical footprinting samples. We are very grateful to Yuriy Chaban and Diamond Light Source for access and support of the Cryo-EM facilities at the UK national electron bio-imaging centre (eBIC), proposal EM-BI24557, funded by the Wellcome Trust, MRC and BBSRC. The initial grid screening was performed in the cryo-EM facility in School of Biological Sciences at the University of Edinburgh, which was set up with funding from the Wellcome Trust (087658/Z/08/Z) and SULSA. We are also grateful to Nathan Cowieson at Diamond Light Source for access and support at the B21 beamline (rapid access proposal SM27169). Overall, MDW’s work is supported by the Wellcome Trust [210493], Medical Research Council (T029471/1), and University of Edinburgh. GD’s work is supported by BBSRC EastBIO [BB/M010996/1]. This work was also supported by the Edinburgh Protein Production Facility (EPPF) and the WCB Bioinformatics Core Facility, both of which receive funding from a core grant (203149) to the Wellcome Centre for Cell Biology at the University of Edinburgh.

## Competing Interests

The authors declare no competing interests.

## Author contributions

MDW conceived the study and supervised the project. GD and MDW designed the experiments (unless otherwise stated), analyzed the data, and wrote the manuscript, with input from the other authors. MRDT, RC, GD, MDW, and HW purified protein and DNA components. Cryo-EM image processing was performed by MDW, GS, and MWT. Model building was performed by MDW, GD, and HB. JW performed DNA footprinting assays. Biochemical assays were performed by MDW, GD, and HW. Phylogenetic analysis and computational protein structure analysis was performed by GD. MNase sequencing data were processed by SW and GD.

## Figures

## Supplementary Figures

**Supplementary Movie 1: overview of Cryo-EM density map and built model** Map coloured according to density attributed to histones (H2A=yellow, H2B=red, H3=blue, H4=green) or DNA (grey), using colour zone tool in ChimeraX.

**Supplementary Movie 2: comparison between *H. sapiens* and *T. brucei* NCP** Overview showing compression and overall oval shape, then zoomed view of SHL6 and SHL2. Created using the morph feature in ChimeraX, moving between *T. brucei* NCP to *H. sapiens* NCP

## Materials and Methods

### Generation of Plasmid Constructs

Plasmids encoding histones from *T. brucei brucei* (protein sequences corresponding to H2A: Tb927.7.2820, H2B: Tb927.10.10480, H3: Tb927.1.2430 and H4: Tb927.5.4170) were received as a gift from the Janzen lab^34^. We refer to the species *T. brucei brucei* as ‘*T. brucei*’ only throughout the study. Histone mutations were made either using site directed mutagenesis or cloning of synthesised double-stranded gBlock fragments containing mutations (Integrated DNA Technologies).

### Histone Protein Purification

Histones were expressed in BL-21 DE3 RIL cells and purified from inclusion bodies essentially as described previously^32–34, 84^ with some modifications. Briefly, for all *H. sapiens* histones and *T. brucei* H2A, H3 and variants, inclusion bodies were extensively washed (50 mM Tris pH7.5, 100 mM NaCl, 1 mM EDTA, 1 mM benzamidine, 5 mM BME), disrupted in DMSO, and resolubilised (7M Guanidine-HCl, 20 mM Tris pH 7.5, 5 mM DTT). They were then dialysed into urea dialysis buffer (7M urea, 100 mM NaCl, 15 mM Tris pH 7.5, 1 mM EDTA, 5 mM BME) for 12h. Dialysis and the cation chromatography step for *T. brucei* histones H2B and H4 were performed at reduced pH 7. Human NCPs were assembled using sequences corresponding to human H2A.1, H2B.1, H3.1 C96S C110A, and H4 and are referred to as *H. sapiens* NCPs throughout.

Histone protein mass was confirmed by 1D intact weight ESI mass spectrometry (SIRCAMs, School of Chemistry, University of Edinburgh) (Supplementary Fig. 2a). Concentrations were determined via absorbance at 280 nm using a Nanodrop One spectrophotometer (Thermo Scientific).

### GST-PFV-GAG Protein Purification

A peptide derived from Prototype Foamy Virus (PFV) GAG (UniProt ID: P14349, aa. 535-550) that binds to the acidic patch^55^ was cloned into a pET His_6_-GST-TEV-LIC plasmid vector and expressed recombinantly in *E. coli* BL21 DE3 RIL cells using overnight induction at 18°C with 0.4 mM IPTG. The protein was then purified using nickel affinity chromatography (HiTrap IMAC HP, Cytiva) (20 mM Tris pH7.5, 400 mM NaCl, 10% glycerol, 2 mM BME, 15-400 mM imidazole gradient) and size exclusion chromatography (HiLoad 16/600 Superdex 200, GE Healthcare) (150 mM NaCl, 5% glycerol, 15 mM HEPES, 2 mM DTT).

### NCP Reconstitution

NCPs were reconstituted essentially as described^32, 33^, with some alterations to improve stability. Briefly, DNA for wrapping all NCPs except for hydroxyl radical footprinting assays was generated by isolating large-scale quantities of the plasmids pUC57 8 x 145bp Widom-601 DNA or 32×147 bp alpha-satellite DNA by multiple rounds of MaxiPrep Kit purifications (Quiagen). The 145 bp fragments were digested and extracted from the plasmid using EcoRV digestion and subsequent PEG and ethanol precipitation steps. For hydroxyl radical footprinting, linker DNA was required to avoid initial undigested signal and the 175bp Widom-601 sequence was used. Fluorescently-tagged DNA was generated by PCR essentially as described^84, 85^:

Fwd primer: 5’TAMRA-ATGGAACACATTGCACAGGATGTAT

Rev primer: 5’6-FAM AATACGCGGCCGCCCTGGAG

Purified octamers were wrapped with the DNA using an 18 h exponential salt reduction gradient. The extent and purity of NCP wrapping was checked by native PAGE and SDS-PAGE analysis (Supplementary Fig. 2b-d). Due to the appearance of a higher molecular weight species in *T. brucei* NCPs, different molar ratios of octamer:DNA (0.4-1.2, were [DNA] = 0.3 mg/ml) were tested to optimize wrapping efficiency (Supplementary Fig. 2b-c). Where necessary, NCPs were purified on HiLoad 16/600 Superdex 200 size exclusion column (GE Healthcare) in 20mM HEPES pH 7.5, 150mM NaCl, 1mM DTT to enrich for NCP only fractions. NCPs were then dialysed for 3h into a customised Storage Buffer (25 mM NaCl, 2.5% glycerol (v/v), 15 mM HEPES pH 7.5, 1 mM DTT), concentrated, and stored at 4°C for maximum of 1 month. For all biochemical experiments, *H. sapiens* and *T. brucei* NCPs were treated identically and processed concurrently.

### Cryo-EM Grid Preparation and Transmission Electron Microsocpy

For cryo-EM grid preparation, NCPs were diluted to a final DNA concentration of 110 μg/ml (DNA concentration) and NaCl concentration of 50 mM. Glutaraldehyde crosslinking agent was added (0.05%) and incubated on ice for 5 minutes. The reaction was quenched with excess ammonium bicarbonate and Tris pH8. NCPs were concentred through a 100 kDa spin concentrator column (Amicon® Devices) and loaded on an HiLoad 16/600 Superdex 200 size exclusion column (GE Healthcare) in 20mM HEPEs pH 7.5, 150mM NaCl, 1mM DTT. Fractions enriched for NCPs were pooled and concentrated (Supplementary Fig. 3c).

Monodispersity of the sample was confirmed by negative staining as described^86^. Briefly, 5 µg/ml NCPs were applied to 300 mesh copper-grids with continuous carbon-film (C267, TAAB) and stained with 2% uranyl acetate for 2 minutes prior to washing. Grids were loaded and imaged in F20 TEM operated at 200kV. Images were collected manually using the EMMENU software (TVIPS) on a TemCam F816 camera (TVIPS) (University of Edinburgh, Transmission EM facility) (Supplementary Fig. 3a).

For single particle cryogenic electron microscopy, 3.5 μl of freshly purified and crosslinked NCPs were applied to glow discharged holey carbon quantifoil R 2/2 grids at a concentration of 2.2 μM. Grids were incubated and blotted at 100% humidity and 4°C in a vitrobot mark IV, prior to vitrification in liquid ethane and storage in liquid nitrogen. Grids were screened for ice quality and a small dataset was collected and processed to 2D classes on a TF20 microscope (University of Edinburgh, Cryo-transmission EM facility). Data collection was then performed on a Titan Krios operated at 300 kV equipped with a Gatan K3, operating in correlated double sampling mode. 4193 Lzw compressed tiff movies were obtained using automated serialEM software^87^ using a pixel size of 0.829 Å and a total dose of 45.7 electrons/Å^2^ (Table 1).

### Cryo-EM Image Processing

All micrographs were motion-corrected using MotionCor2, removing the first frame. CTF parameters were estimated using patch CTF in cryoSPARC^88^ and poor micrographs were discarded. ∼1,000 particles were picked manually and 2D classified to produce templates for template-based picking in cryoSPARC. Two rounds of 2D classification were performed to discard poorly averaged particles and discernible secondary-structure features were pooled. The selected classes were used for ab-initio reconstruction and separated into two ab-initio classes. The best class comprising 306,475 particles were re-extracted with a 316 pixel voxel size and subjected to local CTF refinement and homogeneous refinement with a dynamic mask starting at a resolution of 20 Å and yielding a final map at 3.28 Å resolution. This map was used for all model building and figure preparation. Non-uniform refinement^89^ was performed, removing some noise and yielding a GS-FSC map of 3.22 Å. C2 symmetry was applied in homogenous refinement. These maps were used only to aid map interpretability during model building. Map quality and anisotropy were assessed manually in Chimera^90^ and using 3D-FSC^82^. 3D classification was performed using multiple starting classes using the heterogeneous refinement job in cryoSPARC (Fig. 2d).

### Model Building

The crystal structure of the 145bp Widom-601 DNA was used from PDB: 3LZ0^36^ in the most logically fitting orientation based on Widom 601 DNA asymmetry and best model to map fits. Initial models for *T. brucei* H2A and H2B histones were generated as a dimeric assembly and *T. brucei* H3 and H4 as a tetrameric assembly with glycine linkers in Alphafold2^51^ and docked in the EM map using UCSF ChimeraX^77^. The model was adjusted using Coot^91^ and ISOLDE^92^. Sequences outside of the density were removed manually in Coot and refined using using Phenix real space refinement^93^. Protein geometry was assessed with MolProbity^94^. Model fit was assessed using map-to-model cross correlations^93^ and EMringer^95^. Models have reasonable stereochemistry and are in good agreement with the EM density maps. Figures were prepared in UCSF Chimera and ChimeraX^77^.

### Small Angle X-ray Scattering (SAXS)

SEC-SAXS experiments were performed at Diamond Light Source on the B21 beamline. Freshly prepared *H. sapiens* and *T. brucei* NCPs were loaded on column at 2.4 mg/ml (quantification based on total NCP concentration) and separated by an S200 Increase 3.2 size exclusion column in 20mM HEPEs pH 7.5, 150mM NaCl, and 1mM DTT prior to injection into the beamline and recorded with 3s exposure. Data was reduced and analysed using ScÅtter IV^96^

### Protein Structure Analysis

Structural alignments between *T.* brucei and *H. sapiens* histones were generated using UCSF Chimera or Chimera ^77, 90^. Hydrogen bonding interactions between the histones and DNA in the *T. brucei* structure were calculated using PISA^78^. The overall molecular dipole moments of *T. brucei* and *H. sapiens* histone octamers were predicted using the Protein Dipole Moments Server^80^. The N- and C-terminal tails of *H. sapiens* histones were truncated for this purpose based on the *T. brucei* structure. pKa values of residues in both *T. brucei* and *H. sapiens* H2A-H2B dimers at SHL3.5 were predicted using PROPKA^79^ (Source Data File 1). The surface area of the acidic patch was estimated using PyMOL.

### Protein Sequence and Phylogenetic Analysis

To generate phylogenetic trees for each histone, sequences from 20 different organisms sampling kinetoplastids and other eukaryotes were collected from TriTrypDB^97^ or NCBI Protein, respectively^98^. Multiple sequence alignments were performed with MAFFT^99^ and visualized with Jalview^100^. Maximum likelihood phylogenetic trees were estimated using IQ-TREE^101^, rooted at midpoint, and visualised with iTOL^102^. Heatmaps showing percentage identity to *T. brucei* were generated with iTOL using percentage identity matrices calculated by MUSCLE^103^. Pairwise percentage identities and similarities between *H. sapiens* and *T. brucei* histones were computed using EMBOSS Needle^104^. Isoelectric point (pI) values were obtained from Protparam^105^.

To obtain larger percentage identity matrices, the TriTrypDB^97^ and NCP Protein^98^ databases were mined for histone sequences from 41 organisms (of which 22 were primarily from kinetoplastid reference genomes) and the matrices calculated using MUSCLE. The results were then displayed as correlation maps coloured from 40-100% sequence identity using ggplot2.

### Thermal Denaturation Assays

50 μl reactions with 5 μM NCPs, 20mM HEPES pH 7.5, 150mM NaCl, 1mM EDTA, 1mM DTT, and 5x SYPRO orange (Life Technologies) were set up in a 96-well plate format and heated from 45°C to 95°C with 0.5°C increments on a Biometra TOptical RT-PCR device (excitation/emission λ = 490/580 nm). Relative fluorescence intensity was normalized as (RFU-RFUmin)/(RFUmax-RFUmin). Tetrasome data was normalized from 60°C due to inherent background signal at lower temperatures. Results from three independent experiments, each with two technical repeats were used to calculate T_m_ values for *H. sapiens* and *T. brucei* NCPs (Source Data File 2).

### Salt Stability Assays

250 ng of *T. brucei* and *H. sapiens* NCPs (A260 DNA-based quantification) were incubated at various NaCl concentrations (0.5, 1.0, 1.5, and 2M) for 1h in 10 μl reactions on ice (2.5% glycerol, 15 mM HEPES pH 7.5, 1 mM DTT). After 1h, NaCl concentrations were normalized to 0.15 M and 22 ng of each sample was loaded onto a 5% Tris-glycine polyacrylamide non-denaturing gel. A control sample kept in Storage Buffer (see ‘NCP Reconstitution’) was adjacently loaded.

### Hydroxyl Radical Footprinting

10 µl samples containing 500 ng of fluorescently labelled nucleosomes (5’ 6-FAM labelled reverse strand, 5’-TAMRA labelled forward strand) were set up in reaction buffer (15 mM Hepes pH 7.5, 25 mM NaCl, 1 mM EDTA, 1 mM DTT). 2.5 µl each of 2 mM Ammonium Iron (II) Sulfate/4 mM EDTA, 0.1 M sodium ascorbate, and 0.12% H2O2 were pipetted onto the sides of the reaction tube, mixed together and added to the sample. The reaction was stopped after 4 minutes by the addition of 100 µl STOP buffer (100 mM Tris pH 7.5, 1% glycerol, 325 mM EDTA, 0.1% SDS, 0.1 mg/ml ProteinaseK [Thermo]). The stopped reaction was then incubated for 20 minutes at 56°C to allow ProteinaseK digestion to occur. Fragmented DNA was purified by ethanol precipitation and resupended in 10 µl HiDi Formamide. 0.5 µl of GeneScan 500 LIZ size standard (Thermo) was added as a size marker. The resuspended DNA was run on either a 3130xl Genetic or 3730xl DNA Analyzer, operated using the G5 dye filter set. Peaks were analysed using Thermofisher Connect Microsatellite analysis software. Peak size in base pairs were called by the Global southern method.

### Micrococcal Nuclease (MNase) Digestion Assays

1.2 μg of NCPs (A260 DNA-based quantification) and 7.2 units of MNase were incubated in 70 μl reactions at 37°C (50 mM Tris pH 8.0, 2.5% glycerol, 25 mM NaCl, 5 mM CaCl2, 1.5 mM DTT). The reaction was stopped by mixing 10 μl at relevant timepoints with 5 μl of stop solution (20 mM Tris pH 8.0, 80 mM EDTA, 80 mM EGTA, 0.25% SDS, 0.5 mg/ml Proteinase K). 44 ng of reaction products were then analysed on a non-denaturing, 5% polyacrylamide TBE gel and stained with Diamond DNA stain (Promega). Experiments were performed with *H. sapiens, T. brucei* WT, and *T. brucei Hs*H3 NCPs in triplicate and the disappearance of NCP band intensities were quantified with the BioRad Image Lab software.

### Preparing a Custom DNA Ladder for MNase Products

50 μl restriction digest reactions were set up with 2 μg of 145 bp Widom 601 DNA and HpaII, PmlI, or HpaII + PmlI (1x NEB CutSmart Buffer, 5 U enzyme/μg DNA) for 1h at 37°C. Restriction enzymes were heat inactivated at 80°C for 2 min. Widom 601 171 bp DNA was generated using PCR as described in the NCP reconstitution section. The final ladder comprised five larger fragments (171, 145, 135, 124, and 114 bp). The fragments were mixed at an equal DNA mass ratio and an aliquot of the final mixture (∼20 ng of each fragment) was loaded on a non-denaturing 5% polyacrylamide gel.

### Sequencing MNase Reactions

3 μg of NCPs (A260 DNA-based quantification) and 18 U of MNase were incubated in 150 μl reactions for 30 min at 37°C (50 mM Tris pH 8.0, 2.5% glycerol, 25 mM NaCl, 5 mM CaCl2, 1.5 mM DTT). Reactions were quenched with 75 μl of stop solution (20 mM Tris pH 8.0, 80 mM EDTA, 80 mM EGTA, 0.25% SDS, 0.5 mg/ml Proteinase K). Around 1μg of DNA was isolated from each reaction using a Monarch PCR & DNA Cleanup Kit (elution volume of 15 μl, NEB). The DNA was treated with 5 U of Antarctic Phosphatase (NEB #M0289S) for 1h at 37°C in 20 μl reactions. The phosphatase was heat-inactivated for 2 min at 37°C and the reactions sent for next generation sequencing using the Azenta Amplicon-EZ service (MiSeq 2×250).

### MNase Sequencing Data Processing

168,301 and 132,207 reads were obtained for *H. sapiens* and *T. brucei* respectively, with mean Illumina quality scores > 30. Adapters were trimmed using Cutadapt^106^, aligned to the Widom 601 145 bp sequence using the Burrows-Wheeler Aligner^107^, and filtered for length ≤ 145 bp (0.087% of reads escaped adapter trimming). The final number of mapped reads was 99,212 and 99,050 for *H. sapiens* and *T. brucei* respectively. Read start/end positions extracted from the final dataset are available in Source Data File 3.

### Exonuclease III Assays

Exonuclease assays were preformed essentially as described^108, 109^. Briefly, 2 Units of Exonuclease III (Takara) were added to 1ug of NCPs in ExoIII digestion buffer (50 mM Tris– HCl (pH 8.0), 5 mM MgCl 2, 150mM NaCl and 1 mM DTT). The reaction was incubated at 25°C and samples quenched in stop buffer (20mM Tris pH 8, 200mM NaCl, 0.5% SDS, 25 mM EDTA) at regular intervals. DNA products were deproteinized by digestion with 30 ug of proteinase K followed by ethanol precipitation. Samples were resuspended in HiDi Formamide, prior to running on a denaturing urea 10% polyacrylamide gel and stained with Diamond DNA stain (Promega). The experiment was repeated in triplicate and quantified using BioRad Image Lab software.

### Fluorescence Anisotropy Peptide Binding Assays

NCP-binding FITC-labelled peptides derived from Kaposi’s Sarcoma Herpesvirus Latency Associated Nuclear Antigen (LANA)^54^ and PFV-GAG^55^ were synthesised to > 95% purity by BioMatik, Canada. The LANA peptide with mutations L8A R9A S10A (‘LANA LRS’) and the PFV-GAG peptide with the mutation R540Q (‘PFV-GAG RQ’) were also synthesized and used as non-binding controls. The peptide sequences used are given below:

LANA (UniProt ID: Q9DUM3, aa. 4-23): FITC-Ahx-PGMRLRSGRSTGAPLTRGSC-Amidation

LANA LRS (UniProt ID: Q9DUM3, aa. 4-23): FITC-Ahx-PGMRAAAGRSTGAPLTRGSC-Amidation

PFV-GAG (UniProt ID: P14349, aa. 535-550): FITC-Ahx-GGYNLRPRTYQPQRYGG-Amidation

PFV-GAG RQ (UniProt ID: P14349, aa. 535-550): FITC-Ahx-GGYNLQPRTYQPQRYGG-Amidation

Fluorescence anisotropy assays were performed essentially as described^110^. 50 nM of peptide tracer was incubated with an increasing concentration of *T. brucei* or *H. sapiens* NCPs from 12.5 nM to 2.4 μM (corresponding to 25 nM to 4.8 μM NCP binding sites) in 20mM HEPEs pH7.5, 150mM NaCl, 0.5mM EDTA, 1mM DTT, and 0.05 % Triton X-100. Assays were performed in 25 μl reactions in a 384 well plate and incubated covered at 20°C for 30 minutes. Fluorescence polarization was measured in an M5 multimode plate reader (Molecular Devices), with 480 nm excitation and 540 nm polarised filters (cut-off at 530 nm). Anisotropy (*r*) was calculated as below, where the grating factor (*G*) was approximated to 1, based on FITC alone measurements:

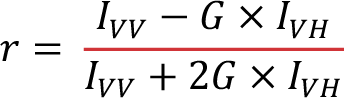

𝐼_*VV*_ = fluorescence intensity of vertically polarized light

𝐼_*VH*_ = fluorescence intensity of horizontally polarized light

The resulting values were background subtracted (no NCPs) and plotted against the number of binding sites on the NCPs (2 protein-binding faces per NCP). The experiment was performed in triplicate (Source Data File 4). The resulting graph was fitted using GraphPad Prism with a non-linear regression, saturation binding curve assuming one site with total and non-specific binding. Binding affinity measurement was estimated using the equation:

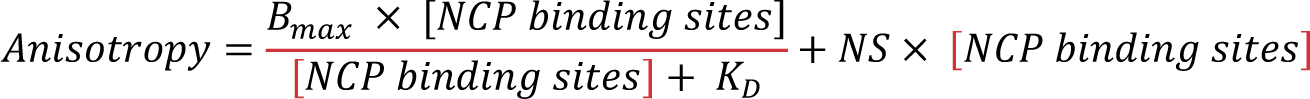

[𝑁𝐶𝑃 𝑏𝑖𝑛𝑑𝑖𝑛𝑔 𝑠𝑖𝑡𝑒𝑠] = molar concentration binding sites in μM (1 on each face of 1 NCP)

𝐵*_max_*_’_ = maximum anisotropy from specific binding

𝐾*_D_* = equilibrium dissociation constant

𝑁𝑆 = slope of non-specific binding (μM^-1^)

### Electrophoretic Mobility Shift Assay for Acidic Patch Binding

30 ng of NCPs (DNA-based quantification) were incubated with different concentrations of the purified GST-His-TEV-tagged PFV GAG (0, 0.2, 0.4, 0.6, 0.8, 1.0, 1.2, 1.4, and 1.6 μM) for 30 min in 15 mM HEPES pH 7.5, 150 mM NaCl, 1 mM DTT, 0.5 mM EDTA, 2.5 % glycerol and 0.05 mg/ml BSA at 4°C. Reaction products (20 ng) were then analysed on a non-denaturing, 5% polyacrylamide TBE gel. Gels were stained with Diamond DNA stain (Promega) and the disappearance of the NCP band quantified using ImageLab. Quantification data can be accessed in Source Data File 4.

---

**Supplementary Figure 1:**
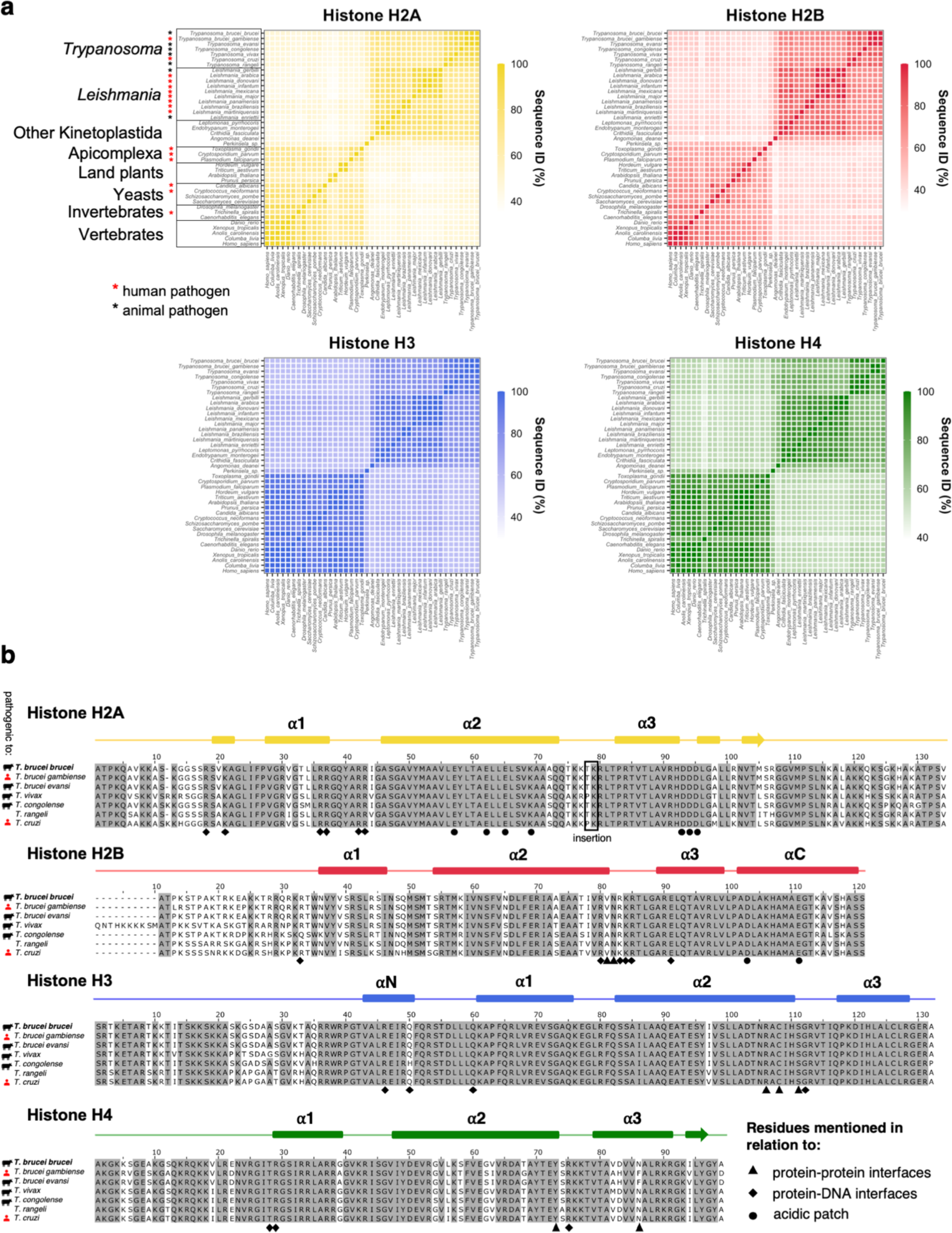
*T. brucei* histone sequences are conserved within kinetoplastids but highly divergent compared to other eukaryotes. **a.** Pairwise sequence identity (ID) matrices of histones from various eukaryotic species. Groups of species that are present in all four matrices are outlined in the panel for histone H2A. Human and animal pathogens are marked with red and black asterisks respectively. **b.** Multiple sequence alignment of histone sequences from *Trypanosoma* species with annotations highlighting residues that are mentioned in relation to histone-histone interfaces, histone-DNA interfaces or the acidic patch in this study. Secondary structure annotations are based on the *T. brucei* NCP model presented in this study.

**Supplementary Figure 2:**
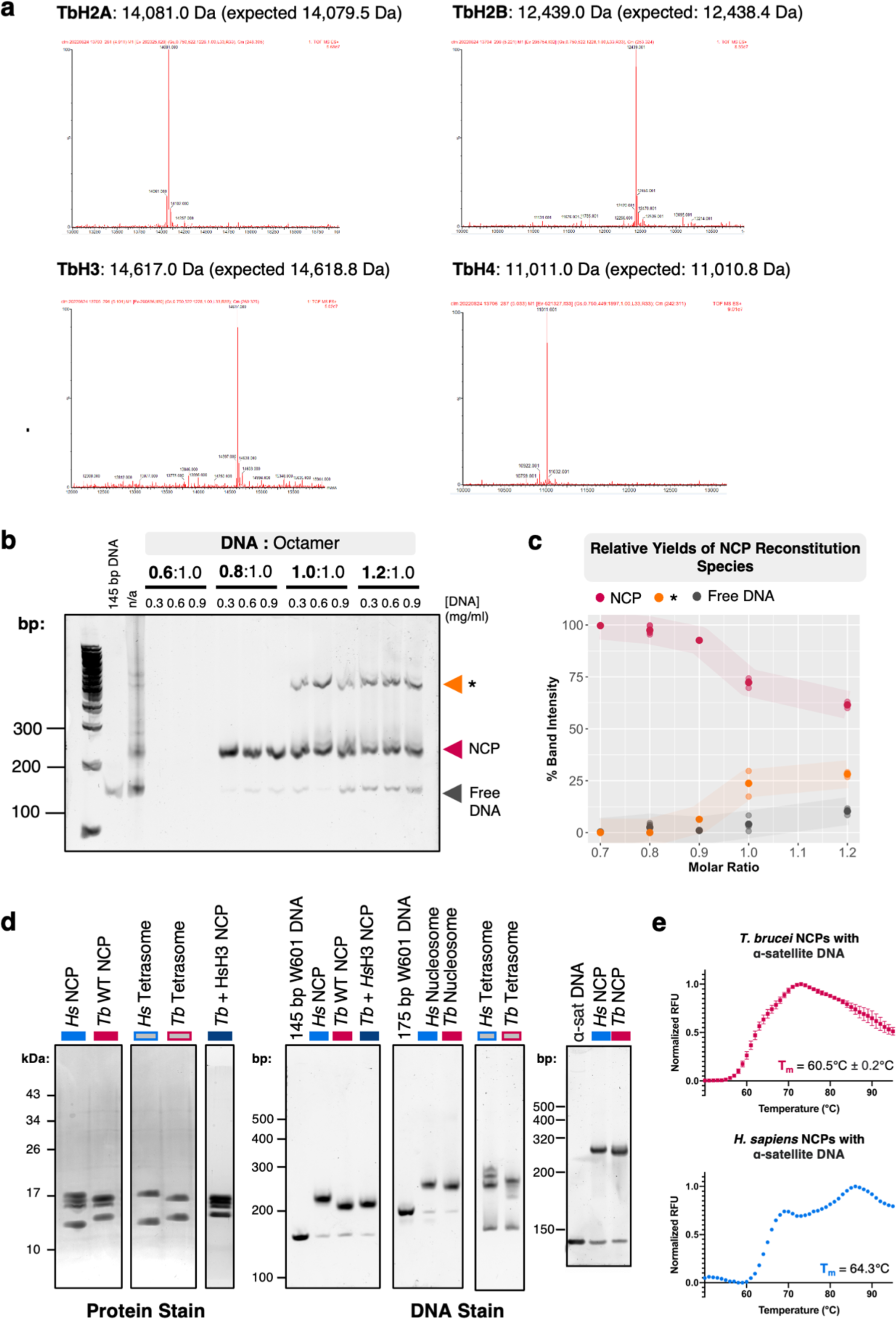
Quality control of *in vitro* reconstituted NCPs and tetrasomes. **a**. 1D Intact mass spectrometry analysis of each *T. brucei* histone showing their deconvolved mass profile. **b**. A range of DNA:octamer molar ratios tested to optimize wrapping of *T. brucei* histone octamers with different concentrations of Widom 601 145 bp DNA (wrapped NCPs = pink, free DNA = grey, unknown higher molecular weight species (‘*’) = orange). **c**. Quantification of relative band intensities from optimization experiments such as **b**. **d**. Gels showing NCPs reconstituted *in vitro* with Widom 601 145 bp DNA and alpha satellite 147 bp DNA, nucleosomes reconstituted with 175 bp FAM-labelled DNA, and H3-H4 tetrasomes consisting reconstituted with 145 bp Widom 601 DNA. On the left, SDS-PAGE gels show equivalent histone distribution. On the right, native polyacrylamide gels show the shift in electrophoretic mobility of DNA when wrapped into NCPs/nucleosomes/tetrasomes. **e**. Thermal denaturation assays of *T. brucei* (top) and *H. sapiens* (bottom) NCPs wrapped with 147 bp alpha-satellite DNA (melting temperatures (T_m_) are indicated).

**Supplementary Figure 3:**
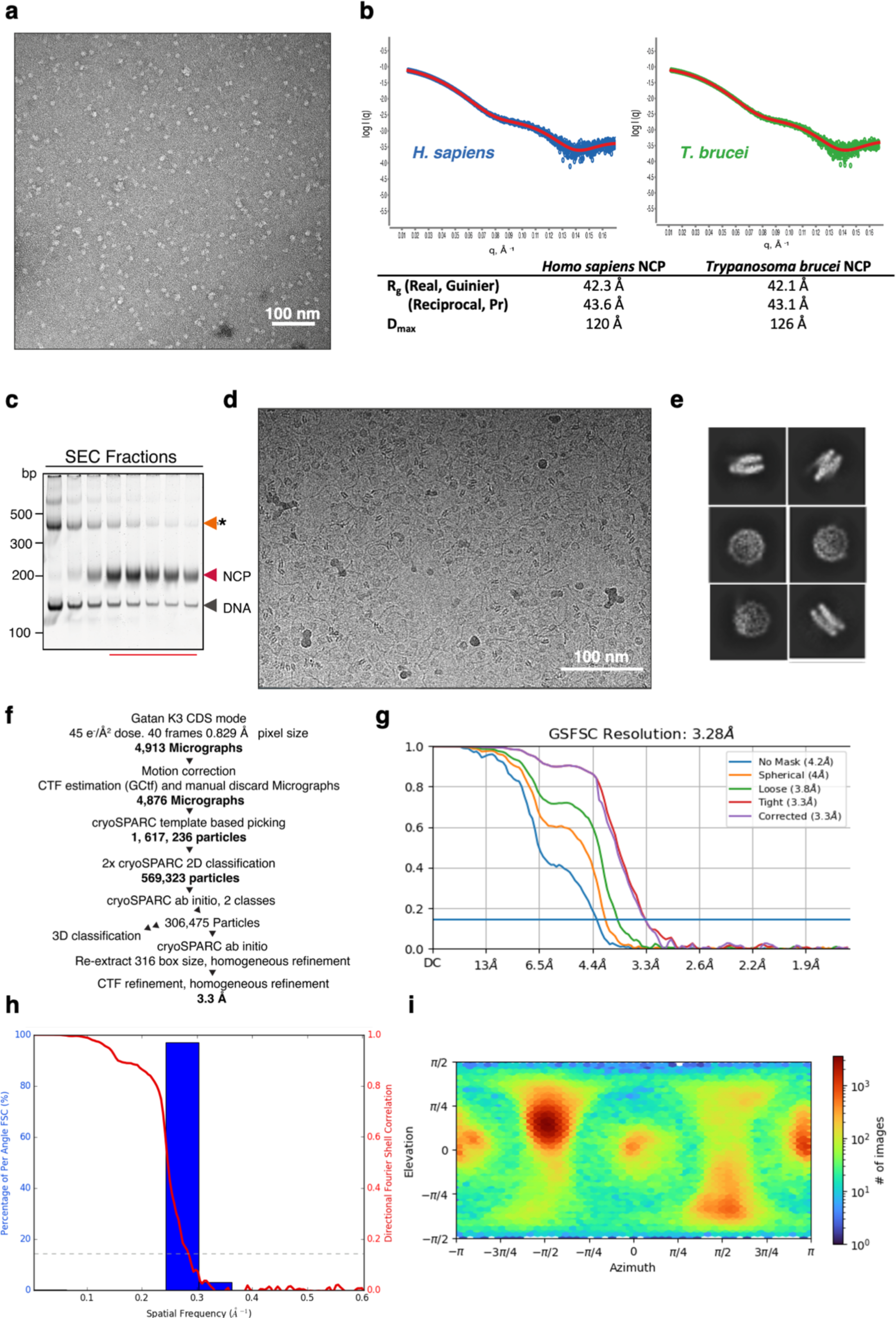
Structural and biophysical characterization of the *T. brucei* NCP. **a**. Representative negative stain EM micrograph of the T. brucei NCP, showing dispersed primarily ∼10 nm top view images. **b**. (top) SAXS scattering profile of and *H. sapiens* (left, blue) and *T. brucei* (green, right) with scattering model fit. (bottom). Table summarizing the radius of gyration (Rg) and maximum dimension (Dmax) calculated from the data using both Porod-Debye (Pr) distribution or normalised Guinier analysis. Rg and Dmax values correlate well suggesting that the averaged size between NCPs is similar. **c.** Native polyacrylamide gel of glutaraldehyde crosslinked NCPs, separated by size exclusion chromatography (SEC) form over crosslinked species, aggregate band (orange) and free DNA (grey). Pooled fractions for cryo-EM grid preparation are indicated with red line. **d**. Representative cryo-EM micrograph of the *T. brucei* NCP. **e**. Examples of 2D class averages obtained during image processing. **f**. flowchart describing cryo-EM image processing pipeline. **g**. Gold-standard Fourier shell correlation (GS-FSC) curve for final map, including unmasked and masked curves.The blue line corresponds to 0.143 threshold. **h.** Three dimensional FSC^82^ for final map showing global FSC curve (red) and overlap of histogram of directional FSC with the major peak correlating with the global resolution estimate. **i.** Euler angle distribution plot of all particles used in the final map. Despite some preferred orientations (red on heat map) no anisotropy in model was observed (h).

**Supplementary Figure 4:**
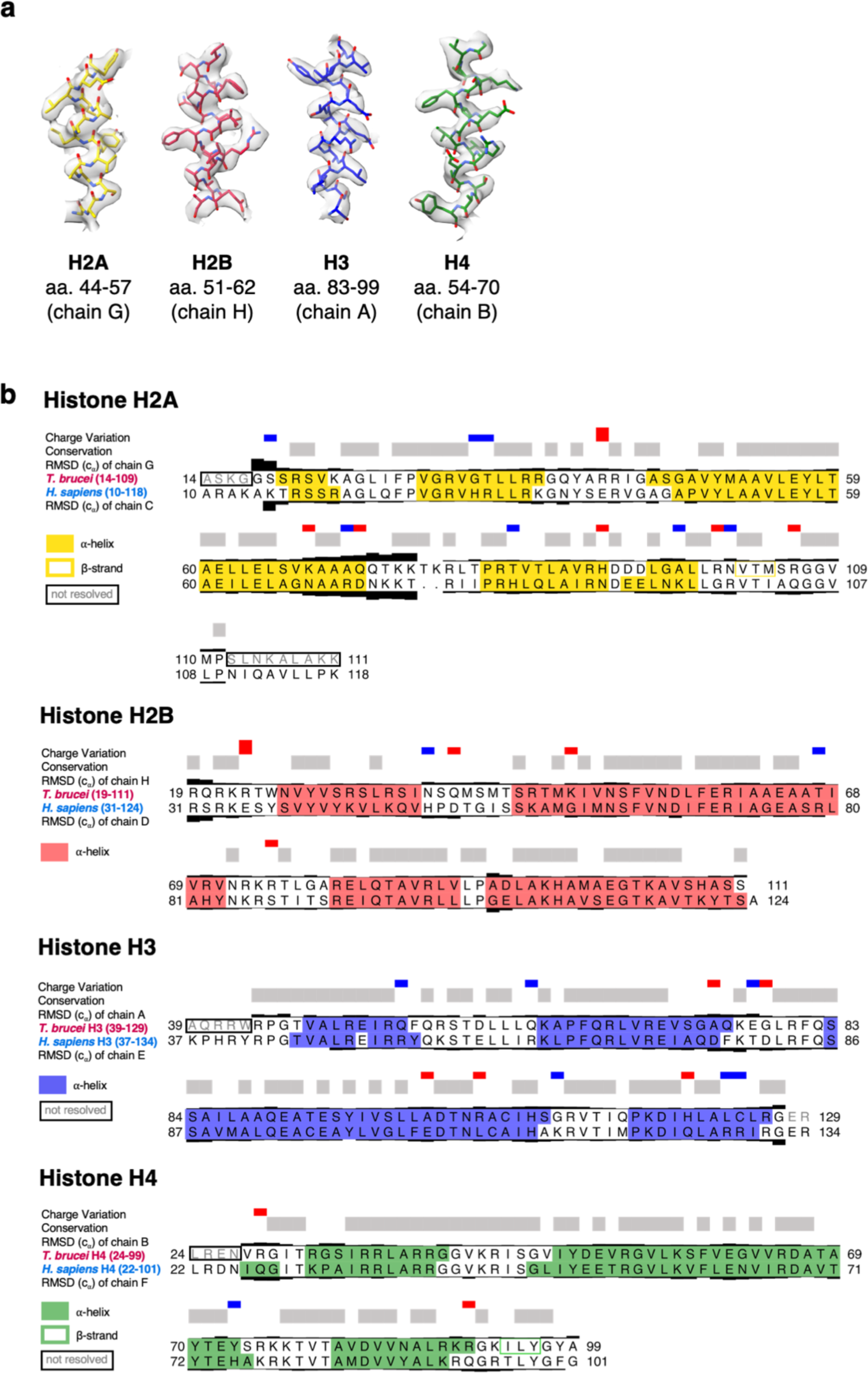
Histone secondary structure and alignment in the *T. brucei* NCP. **a.** Representative images showing the EM density and model building for each *T. brucei* histone**. b.** Pairwise structural alignment of histones from *T. brucei* (our model) and *H. sapiens* (PDB: 7XD1)^42^. **Charge variation** highlights changes in *H. sapiens* sequences compared to *T. brucei*, where blue = change to a positively charged residue, red = change to a negatively charged residue, and the height of the bar represents the magnitude of the change. **Conserved residues** are indicated in grey and the **RMSD values** of c_⍺_ atoms in both chains are shown as black bars, where a larger bar indicates higher RMSD.

**Supplementary Figure 5:**
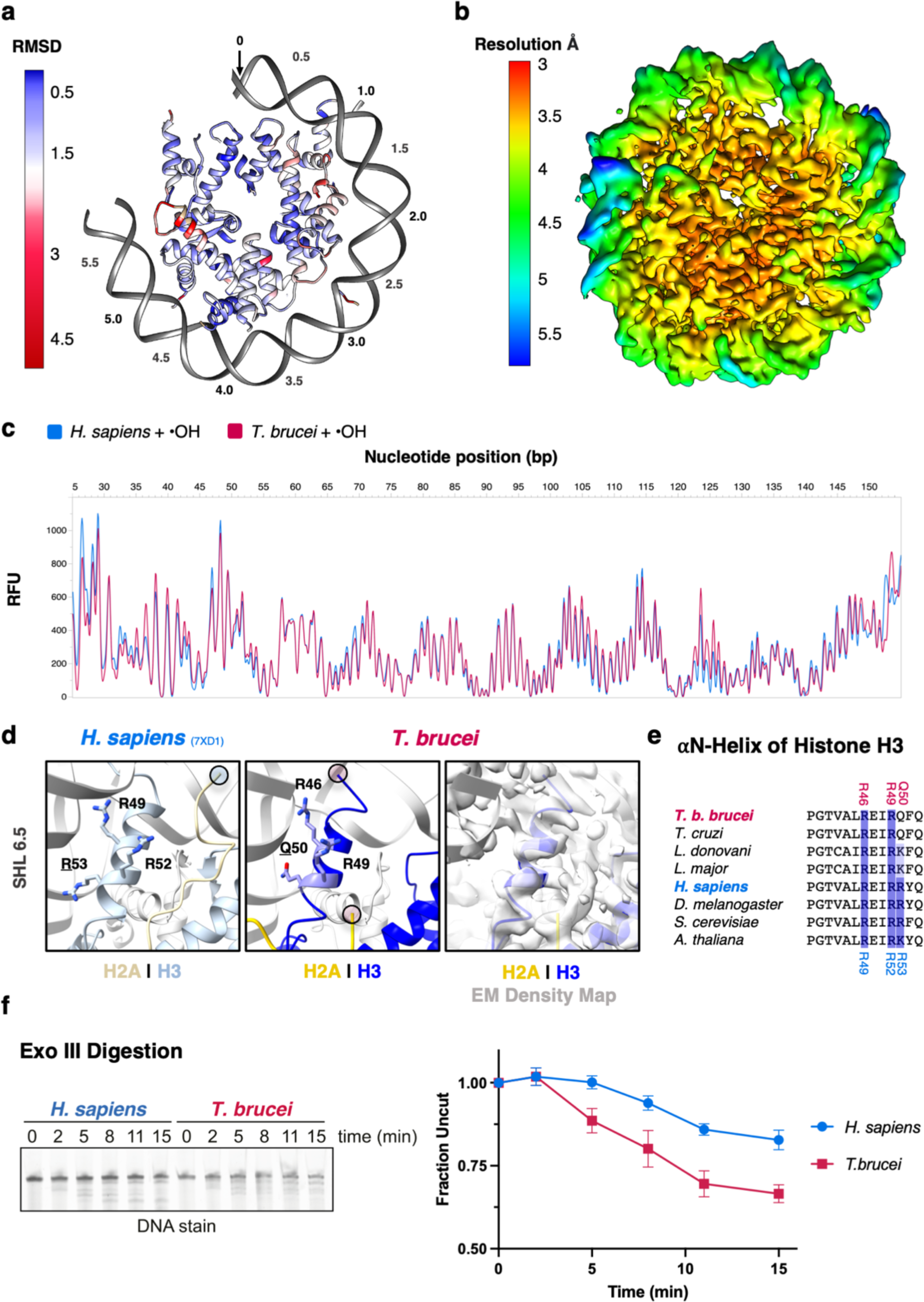
Weakened DNA binding in the *T. brucei* NCP occurs due to alteration of local protein-DNA contacts. **a**. Lateral view *of T. brucei* NCP model, histones coloured according to root mean square deviation (RMSD) between C_⍺_ atoms in this structure and the structure of the *X. laevis* NCP (PDB: 3LZ0)^36^. Only one side of the NCP is shown for clarity. **b**. Cryo-EM density *of T. brucei* NCP coloured according to local resolution, estimated in CryoSPARC. The core of the NCP is highly ordered while the DNA ends are flexible due trypanosome-specific histone sequence alterations. **c**. Hydroxyl radical (•OH) footprinting assay on *H. sapiens* and *T. brucei* nucleosomes wrapped with Widom 601 175 bp FAM-labelled DNA (detectable fragment size range 25 bp – 155 bp). ∼10 bp periodicity of protected residues corresponding to histone-DNA interactions can be seen for both *T. brucei* and *H. sapiens* NCPs, suggesting that overall DNA register is maintained. **d**. Comparison of the N-terminal end of H3 and the C-terminal end of H2A at SHL6 in *H. sapiens* (left) and *T. brucei* (right), including weaker EM density (far right). Circles indicate the end of buildable density on histone tails. **e**. Multiple sequence alignment of the ⍺N helix of histone H3 with DNA. Residues that were previously identified as crucial for contacting DNA^14, 44^ are highlighted. **f**. Denaturing urea gel (left) and quantification of the disappearance of the 145 bp DNA (right) from an Exonuclease III digestion assay of intact NCPs at indicated timepoints.

**Supplementary Figure 6:**
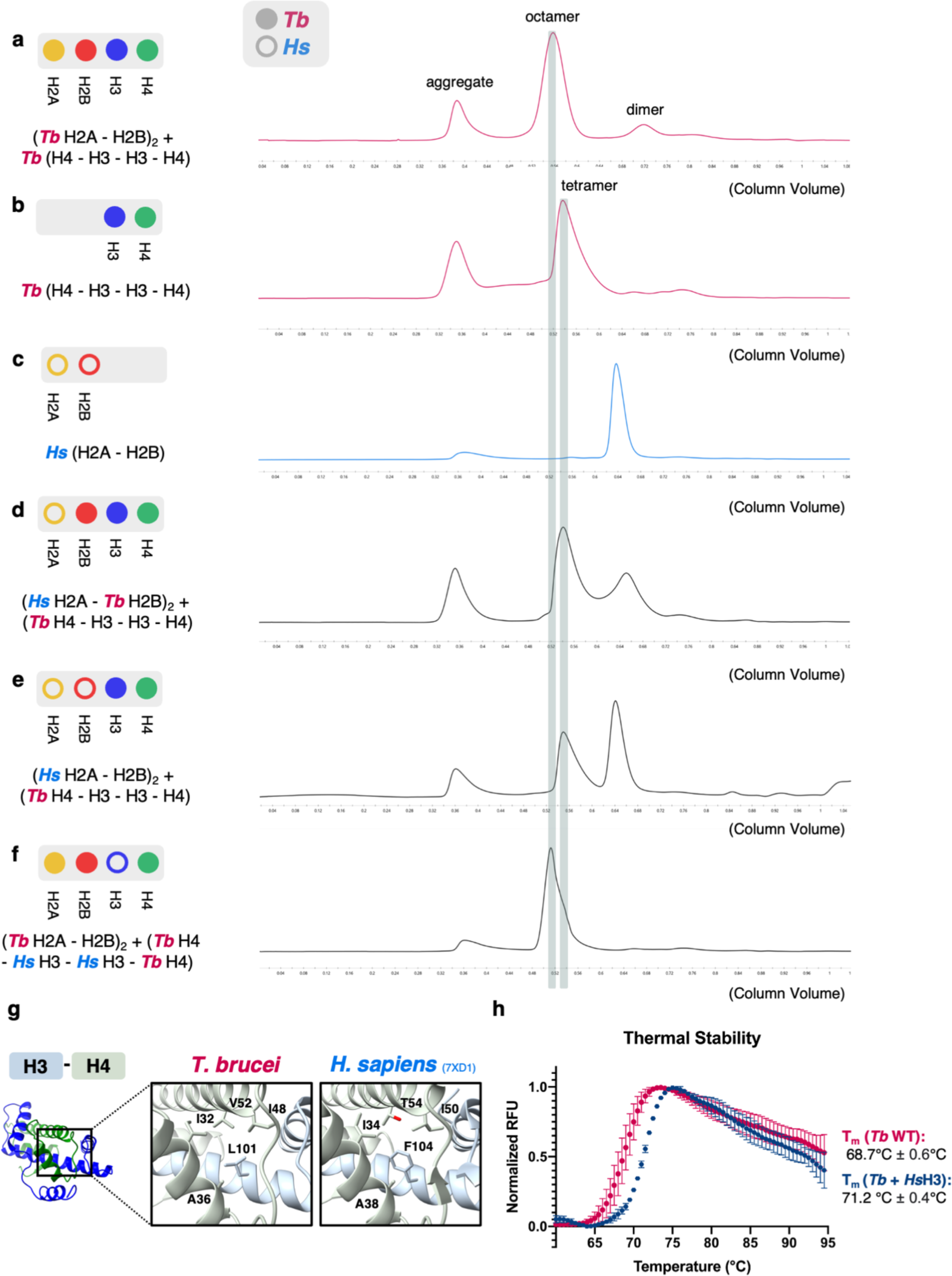
Divergent interfaces in the *T. brucei* NCP dictate the formation and stability of chimeric histone assemblies. Size exclusion chromatograms of various histone complexes after the refolding step of octamer/tetramer/dimer assembly showing lack of assembly for some chimeric octamers. Expected elution volume for the different species is highlighted. **a.** *T. brucei* octamers (used for “Tb WT” NCPs); **b.** *Tb* H3-H4 tetramers; **c.** *Hs* H2A-H2B dimers; **d.** *Tb* H2B, H3, and H4 *Hs* H2A; **e.** *Tb* H3-H4 tetramers with *Hs* H2A-H2B; **f.** *Tb* H2A, H2B, and H4 with *Hs* H3 (used for *“T. brucei* + *Hs*H3” NCPs). **g.** Comparison of the H3-H4 interface in *T. brucei* and *H. sapiens.* **h.** Comparison of thermal denaturation curves from *T. brucei* WT and *T. brucei* + *Hs*H3 NCPs (the only chimeric NCP that could be generated).

**Supplementary Figure 7:**
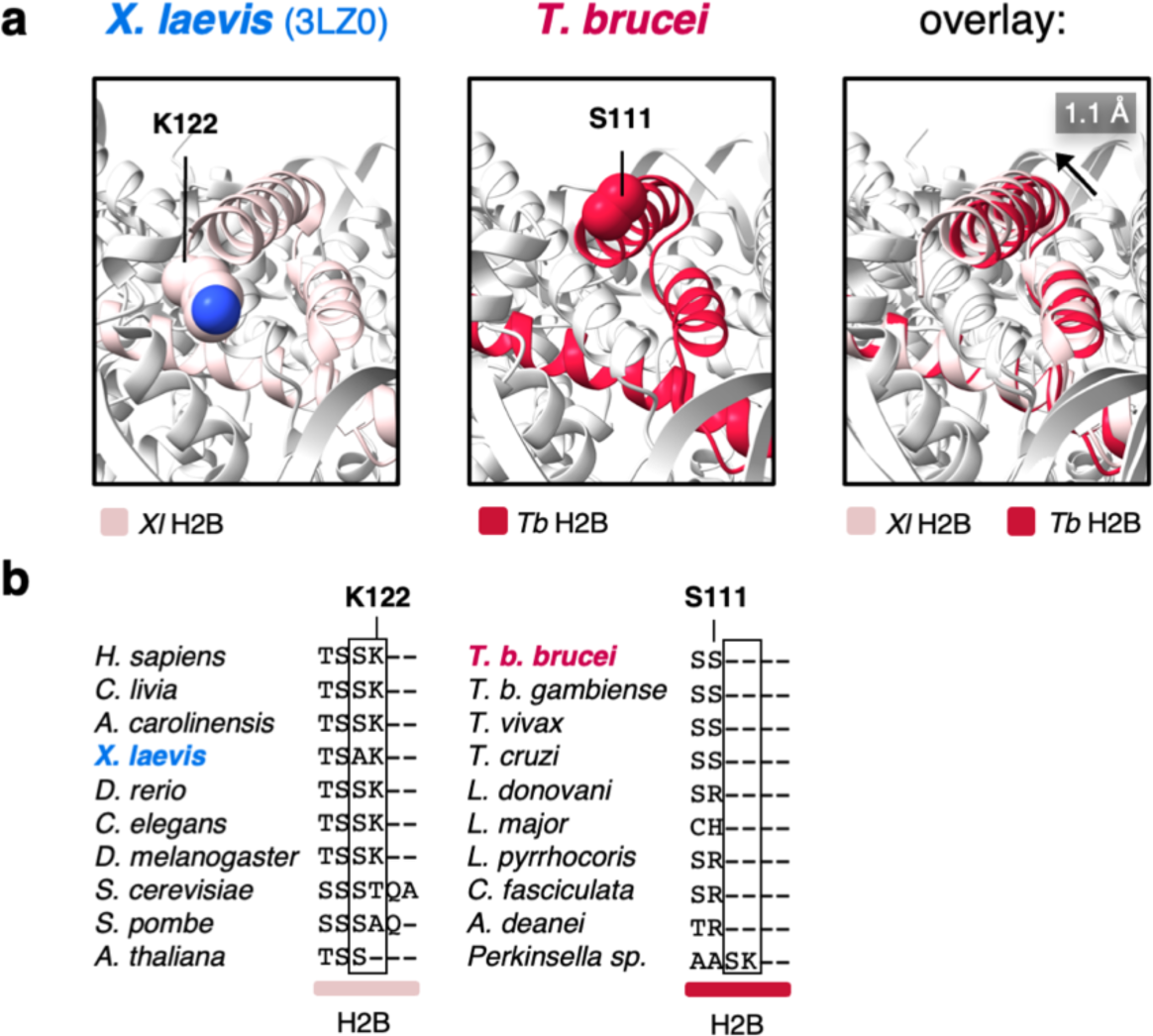
DNA contacts at SHL4.5 are reduced due to altered histone structure. **a.** Comparison of the C-terminal helix of histone H2B in *X. laevis* (left)^36^ and *T. brucei* (right), highlighting the lack of a terminal lysine residue and altered packing in the *T. brucei* NCP (far right). **b.** Multiple sequence alignment of the end of C-terminal helix of H2B. H2B is shorter by two residues in kinetoplastids compared to most of the other eukaryotes.

**Supplementary Figure 8:**
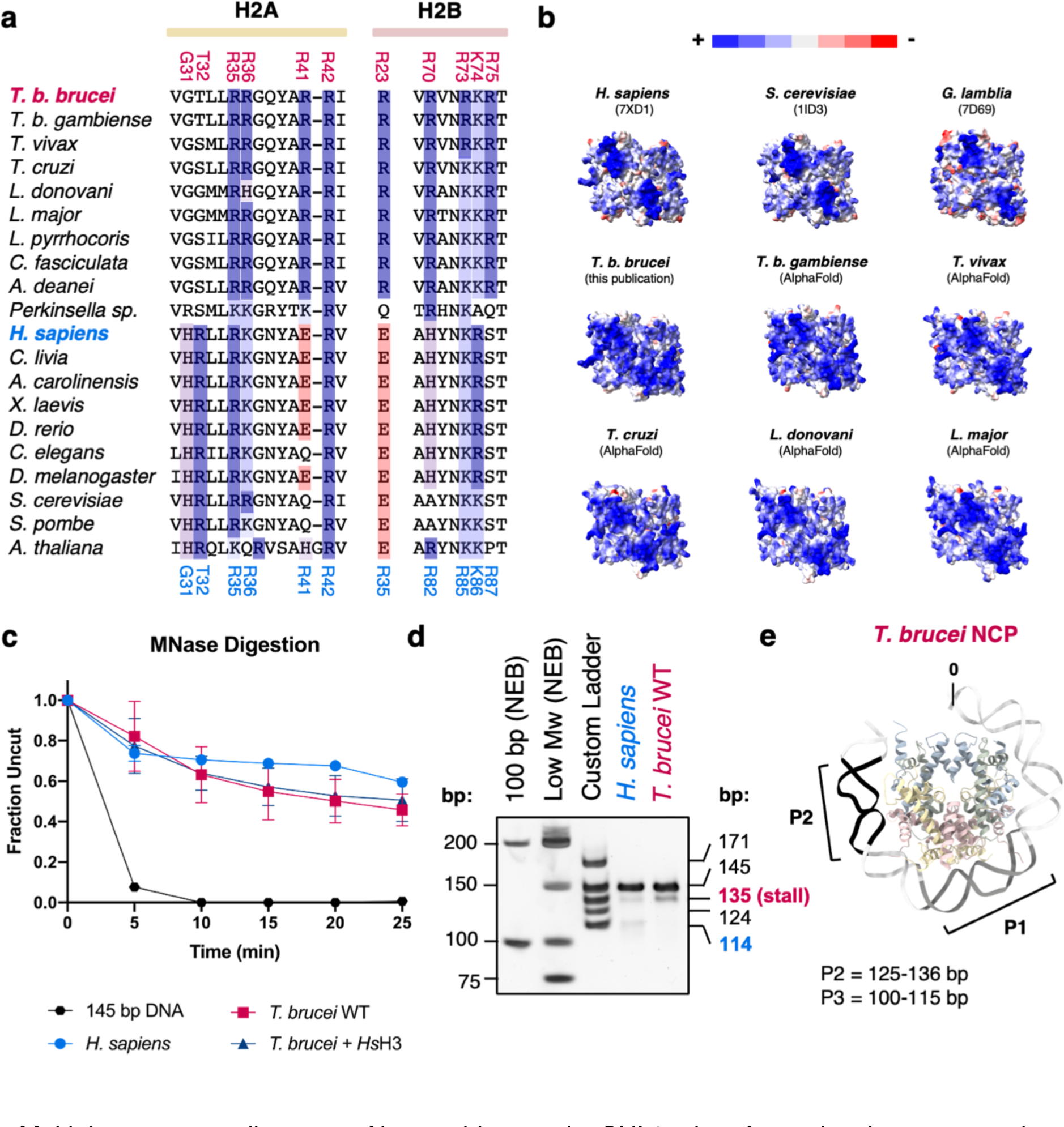
A cluster of positively charged residues drives increased DNA binding at SHL3.5 in the *T. brucei* NCP. **a**. Multiple sequence alignment of key residues at the SHL3.5 interface, showing conservation of extra basic residues predicted in this region. **b**. Comparison of the interface at SHL3.5 in histone octamers from multiple species. Models were either created using AlphaFold2^51^ or taken from previously published structures and colored by surface electrostatics. **c.** Quantification of loss of the full-length band from MNase digestion assays (representative images in Fig 5d) from three independent experiments **d.** Native polyacrylamide gel of MNase digestion products from *H. sapiens* and *T. brucei* NCPs after 30 min next to multiple DNA ladders to map the *T. brucei*-specific stall point. **e.** MNase ‘stall points’ at Peaks 1 and 2 (P1, P2 from Fig. 5e-f in the main text) mapped onto the structure of the *T. brucei* NCP. Peak 1 is the major stall point and digestion occurs predominantly from one side of the NCP.

**Supplementary Figure 9:**
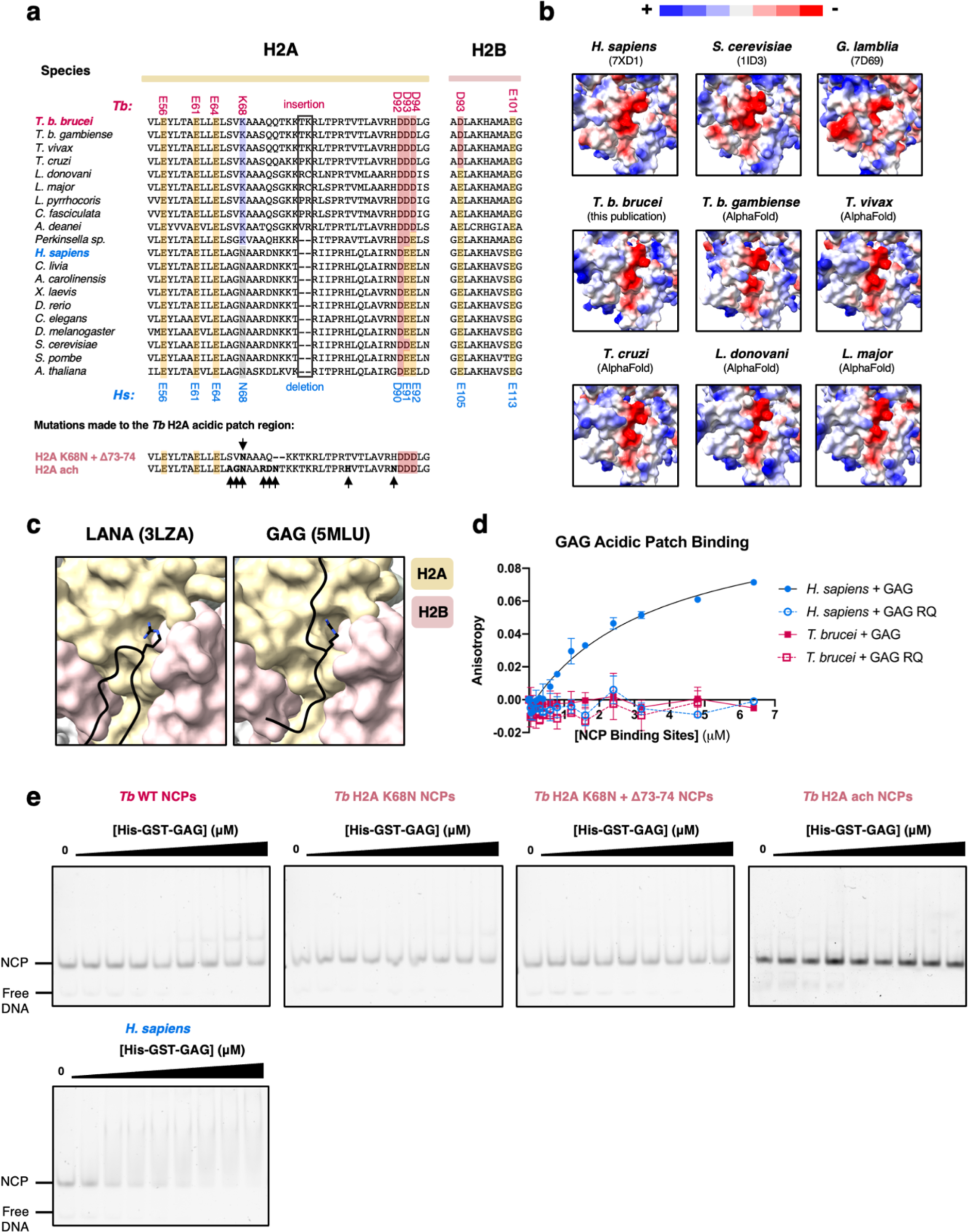
The kinetoplastid acidic patch is highly divergent and extensive mutagenesis of the *T. brucei* acidic patch does not rescue binding to known interactors. **a.** Extended multiple sequence alignment of acidic patch regions in H2A and H2B (see Fig.6a). *Tb* H2A sequences containing ‘humanizing’ mutations that were used in part **e.** are shown below and indicated with black arrows. In **‘H2A K68N + Δ73-74’**, *Tb* H2A-Lys68 is mutated to *Hs* H2A*-*Asn68 and two residues are deleted to remove the kinetoplastid-specific insertion. In **‘H2A-ach’**, multiple mutations are made to residues surrounding the acidic patch by comparing sequence conservation in kinetoplastids and other organisms (S66A + V67G + K68N + A71R + Q72D + Q73N + T84H + H91N). **b.** Comparison of the acidic patch region in histone octamers from various species shown with surface electrostatics, either predicted using AlphaFold2^51^, from *T. brucei* (this study, see Fig. 6), or from previously published NCP structures^10, 42, 56^. **c.** Differential acidic patch binding modes of LANA^54^ and GAG^55^ peptides based on published crystal structures. **d.** Fluorescence polarization assay with a FITC-tagged GAG peptide and a mutated, non-binding GAG peptide (‘GAG RQ’) vs. *H. sapiens* and *T. brucei* NCPs. **e.** Electrophoretic mobility shift assays with His-GST-tagged GAG peptide incubated at increasing concentrations with various NCPs including *H. sapiens*, *T. brucei* WT, *T. brucei Tb* H2A K68N and *Tb* H2A ach ([His-GST-GAG] (μM): 0, 0.2, 0.4, 0.6, 0.8, 1.0, 1.2, 1.4, 1.6; K_D_ ∼ 3.5 μM).

**Supplementary Figure 10:**
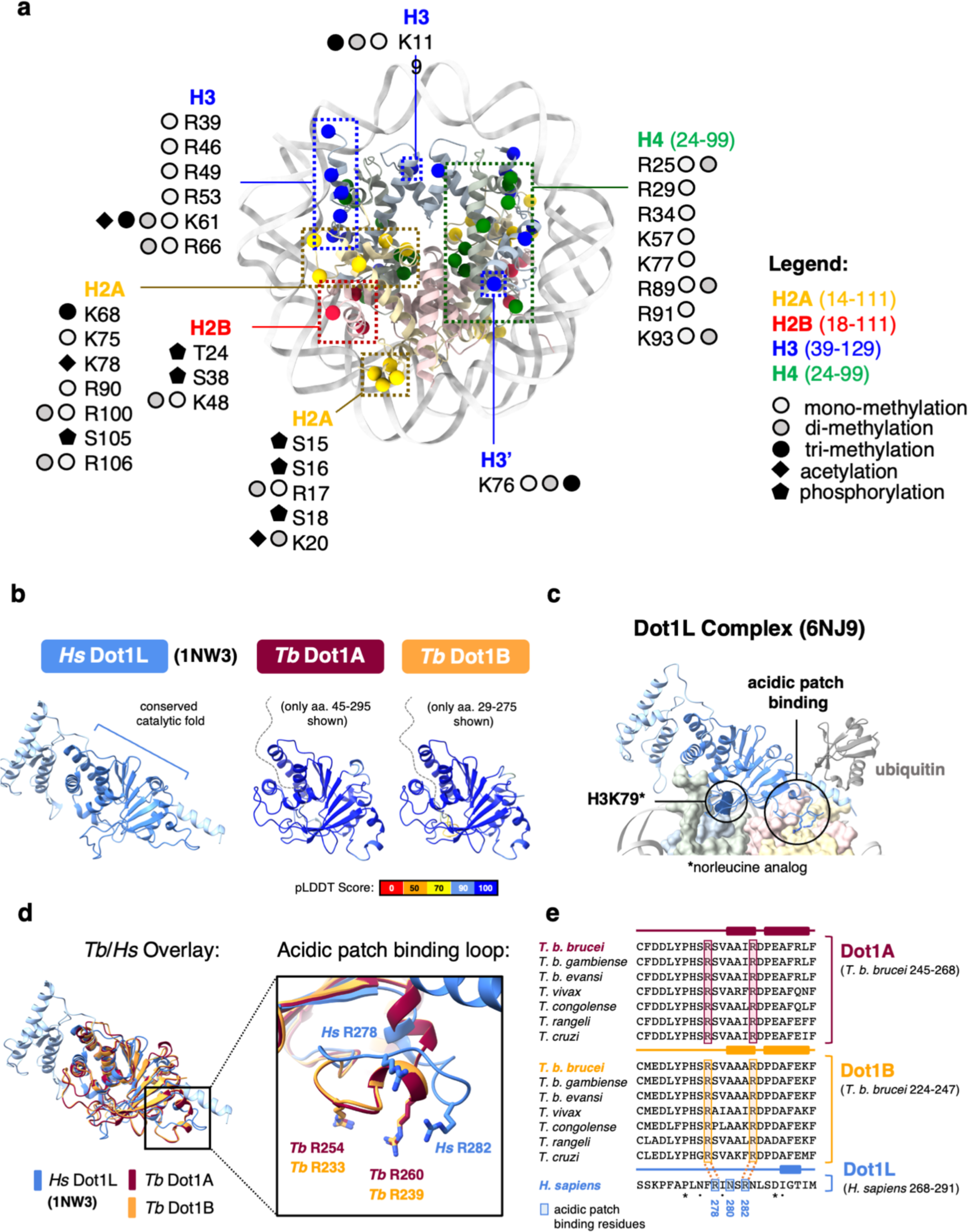
The surface of *T. brucei* NCP likely serves as a dynamic interaction site for chromatin binders. **a.** Previously identified histone post-translational modifications^24, 26^ mapped onto the structure of the *T. brucei* NCP. **b**. Conservation of the catalytic fold of *H. sapiens* Dot1L^83^ and *T. brucei* Dot1A/Dot1A predicted by AlphaFold2^51^. Predicted structures are coloured by their pLDDT confidence score. **c**. *H. sapiens* Dot1L binding to the nucleosome by contacting its substrate H3-Lys79, the acidic patch, and ubiquitin.^73^ Histone ubiquitylation does not seem to be conserved in *T. brucei*, and neither is the ubiquitin interacting region. **d**. Overlay of *H. sapiens* Dot1L and *T. brucei* Dot1A/B and a magnified view of the acidic patch binding loop characterized in *H. sapiens*^73, 83^ **e**. Multiple sequence alignment of the putative acidic patch binding region in Dot1A/B sequences from the *Trypanosoma spp.* This loop is highly divergent compared to the sequence of the *H. sapiens* Dot1L region, but conserved in the *Trypanosoma spp.* (asterisk = identical, dot = similar amino acid). Residues in Dot1L that were shown to bind to the acidic patch (R287, N280, and R282)^71, 73^ are highlighted in blue. The potential arginine equivalents in *T. brucei* Dot1A and Dot1B are highlighted in pink and orange, respectively. The asparagine is lacking and is replaced by an alanine in a cluster of small hydrophobic residues that appear to increase the alpha-helical propensity of the loop. Secondary structure elements for each alignment are shown above. These alterations are consistent with the extensive structural divergence of the trypanosome acidic patch.

## Notes

### Competing Interest Statement

The authors have declared no competing interest.

